# Outer Membrane Vesicles from the gut microbiome contribute to tumor immunity by eliciting cross-reactive T cells

**DOI:** 10.1101/2021.11.05.467432

**Authors:** M. Tomasi, E. Caproni, M. Benedet, I. Zanella, S. Giorgetta, M. Dalsass, E. König, A. Gagliardi, L. Fantappiè, A. Berti, S. Tamburini, L. Croia, G. Di Lascio, E. Bellini, S. Valensin, G. Licata, G. Sebastiani, F. Dotta, F. Armanini, F. Cumbo, F. Asnicar, A. Blanco-Míguez, E. Ruggiero, N. Segata, G. Grandi, A. Grandi

**Author notes:** Corresponding authors: Guido Grandi, Professor of Microbiology and Clinical Microbiology, University of Trento, Trento, Italy, Alberto Grandi, PhD, Toscana Life Sciences Foundation, Siena, Italy. These authors equally contributed to this work (M. Tomasi, E. Caproni, M. Benedet).

## Abstract

The gut microbiome plays a key role in cancer immunity. One proposed mechanism is through the elicitation of T cells, which incidentally recognize neo-epitopes arising from cancer mutations (“molecular mimicry (MM)” hypothesis). To support MM, *Escherichia coli* Nissle was engineered with the SIINFEKL epitope (OVA) and orally administered to C57BL/6 mice. The treatment elicited OVA-specific CD8^+^ T cells in the *lamina propria* and inhibited the growth of OVA-B16F10 tumors. Importantly, the administration of Outer Membrane Vesicles (OMVs) engineered with different T cell epitopes elicited epitope-specific T cells and inhibited tumor growth. Microbiome shotgun sequencing and TCR sequencing provided evidence that cross-reacting T cells were induced at the mucosal level and subsequently reached the tumor site. Overall, our data support the role of MM in tumor immunity, assign a new role to OMVs and pave the way to new probiotics/OMV-based anti-cancer immunotherapies.

## Introduction

The gut microbiome plays a fundamental role in cancer immunity and in determining the efficacy of cancer immunotherapy (Zitvogel et al., 2016). A recent epidemiological study has shown that antibiotic-associated dysbiosis can enhance the frequency of certain cancers, including lung, prostate and bladder cancers (Zitvogel et al., 2016, 2018). Furthermore, it was shown that C57BL/6 germ-free or microbiome-depleted mice respond poorly to PD-1/PD-L1 therapy, while the anti-tumor activity of checkpoint inhibitors is potentiated when *Bifidobacterium* species are administered by oral gavage after tumor challenge (Sivan et al., 2015). Moreover, transplantation of fecal microbiome from patients responding to PD-L1 therapy, but not from non-responders, improves the efficacy of checkpoint inhibitors both in animal models and in melanoma patients (Davar et al., 2021; Routy et al., 2018a; Zitvogel et al., 2018). Finally, retrospective analyses in human patients under PD-1/PD-L1 therapy show the deleterious effect of antibiotics administered during the monoclonal antibody treatment (Zitvogel et al., 2016).

The mechanisms through which the gut microbiome influences cancer immunity are poorly defined. Three non-mutually exclusive mechanisms have been proposed. First, the gut microbiome has been shown to release metabolites, such as polyamine, vitamin B16 and short-chain fatty acids, which mediate systemic effect on the host immunity (Pietrocola et al., 2016). A second mechanism envisages a long distance adjuvant effect, which the microbiome exerts by releasing products and cytokines (Cook et al., 2020; Daillère et al., 2016; Sivan et al., 2015; Vétizou et al., 2015; Zitvogel et al., 2016). The third mechanism assumes that gut microbiome antigens are continuously processed by resident DCs, which in turn induce epitope-specific T cells. This T cell population mainly resides in the intestinal epithelium (intraepithelial lymphocytes (IELs)) and in the *lamina propria*, but can eventually disseminate systemically and reach organs and tumors (Routy et al., 2018a). Considering the abundance and diversity of microbial immunogenic epitopes, it is conceivable to believe that some of them induce T cells capable of recognizing homologous neo-epitopes arising from cancer mutations (“molecular mimicry (MM)” hypothesis). The MM hypothesis is particularly attractive since it would assign a previously unpredicted specificity to the anti-tumor activity of the gut microbiome.

The experimental evidence supporting the role of cross-reactive epitopes is still limited. It has been shown that the rare (<2%) long term (>10 years) survivors of pancreatic cancer carry infiltrating cytotoxic T cells specific for a MUC16 neo-epitope, which cross-react with pathogen-associated epitopes (Balachandran et al., 2017). Moreover, bioinformatics analysis of the gut microbiome has revealed the existence of several microbiome antigens with high homology to known immunogenic T cell epitopes of bacterial, viral, and allergic antigens. This has led to propose the existence of microbiome “tolerogenic” and “inflammatory” epitopes which can dampen or increase the immunogenicity toward the homologous antigen-specific T cell epitopes (Bresciani et al., 2016; Pro et al., 2018). More recently, it has been proposed that mimic peptides from commensal bacteria can promote inflammatory cardiomyopathy in genetically susceptible individuals, leading to myocarditis and lethal heart disease (Gil-Cruz et al., 2019). Moreover, *Bifidobacterium breve* was shown to carry a T cell epitope, which cross-reacts with a model neo-antigen present in B16.SIY melanoma cell line and that the presence of *B. breve* in the mouse intestine reduced the growth of B16.SIY tumors in C57BL/6 mice (Bessell et al., 2020). Finally, mice bearing the tail length tape measure protein (TMP) found in the genome of a *Enterococcus hirae* bacteriophage mounted a TMP-specific CD8^+^ T cell response, which improved PD-1 immunotherapy (Fluckiger et al., 2020).

In an attempt to directly demonstrate that MM can influence cancer immunity, one of our strategies is to artificially introduce cancer-specific epitopes in commensal bacteria and see whether the presence of the engineered bacteria in the mouse intestine could induce epitope-specific T cell responses and could influence the development of tumors expressing such epitopes (Figure 1A).

**Figure 1.**
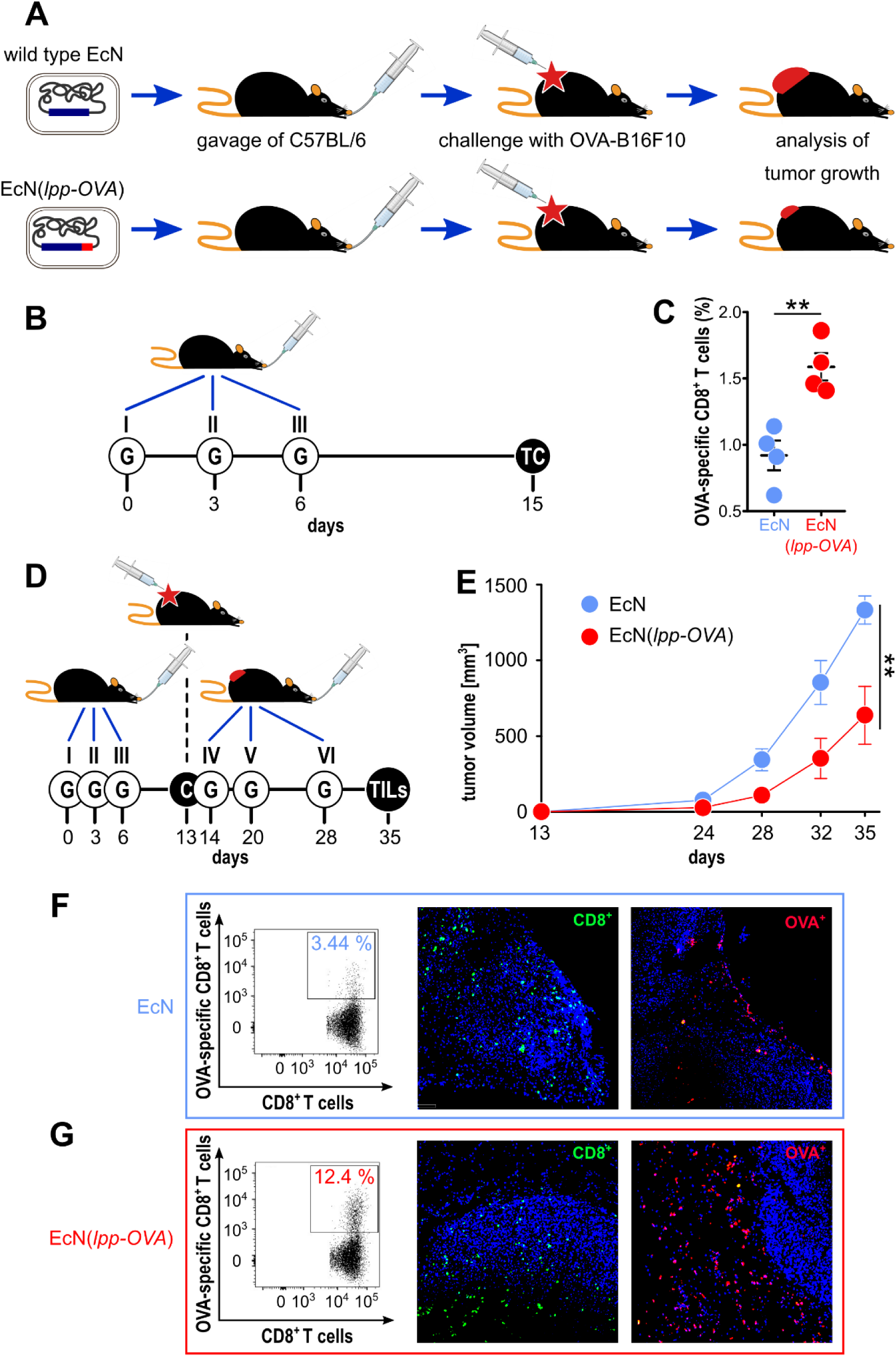
Testing the role of molecular mimicry in tumor inhibition by oral administration of engineered probiotic bacteria. (**A**) *Schematic representation of the experimental strategy used to support the role of “molecular mimicry” in tumor inhibition. E. coli Nissle* was engineered with the OVA CD8^+^ T cell epitope and the strain, or w.t. *E. coli Nissle*, were given to C57BL/6 mice by oral gavage. Animals were subsequently challenged with OVA-B16F10 cells and tumor growth was followed over time. (**B**) *Experimental setup for the analysis of OVA-specific T cells in the lamina propria*. 10^9^ CFU of EcN and EcN(*lpp-OVA*) were given to C57BL/6 mice three times at three day intervals by gavage (“G”). One week after the last gavage, T cells (“TC”) were isolated from the *lamina propria* and OVA-specific CD8^+^ T cells were analyzed by flow cytometry. (**C**) *Flow cytometry analysis of OVA-specific CD8*^*+*^ *T cells in lamina propria* – 1.5×10^6^ cells were extracted from the *lamina propria* of C57BL/6 mice treated with EcN (blue) and EcN(*lpp-OVA*) (red) as described in (B). The frequency of OVA-specific CD8^+^ T cells was measured by using OVA_257-264_ Dextramer-PE. (**D**) *Experimental setup to test tumor inhibition of by oral administration of EcN(lpp-OVA)*. EcN and EcN(*lpp-OVA*) were given at three day intervals to C57BL/6 mice by oral gavage (“G”). One week after the third gavage, mice were challenged (“C”) with 2.8×10^5^ OVA-B16F10 cells followed by three additional gavages. Tumor growth was followed over a period of 23 days and at the end of the experiment tumor infiltrating T cells (TILs) were analyzed. (**E**) *Analysis of tumor inhibition by EcN(lpp-OVA)*. Animals were treated as depicted in D, and tumor volumes were measured over time. Animals were sacrificed when tumors reach a volume of 1.500 mm^3.^ Statistical analysis was performed using Student’s t-test (two-tailed). ^**^ P ≤ 0.01. (**F-G**) *Analysis of Tumor Infiltrating Lymphocytes (TILs) by flow cytometry and immunohistochemistry*. At the end of the experiment depicted in D, tumors were collected dissociated by enzymatic and mechanical treatment and OVA-specific CD8^+^ T cells were analyzed by flow cytometry. The figure reports the analysis of OVA-specific CD8^+^ T cells from one tumor from mice treated with EcN and EcN(*lpp-OVA*) (total CD8^+^ T cells from EcN tumor = 9400/live cells; total CD8^+^ T cells from EcN(*lpp-OVA*) tumor = 8200/live cells). In addition, tumors were fixed and frozen sections were analyzed for OVA-specific CD8^+^ T cells detection using OVA_257-264_ Dextramer labeled with pycoerythryn (red stained cells in panels F and G). Total CD8^+^ T cells (green stained cells) were also visualized by fixing frozen sections in acetone followed by incubation with anti-mouse CD8 rat monoclonal antibody and goat anti-rat Alexa-Fluor 488 conjugate antibody.

## Results

### Oral administration of EcN engineered with a cancer epitope elicit cancer-specific T cells and inhibits tumor growth

The human probiotic *E. coli* Nissle 1917 (Lasaro et al., 2014; Sonnenborn, 2016) was engineered by fusing the SIINFEKL epitope (OVA), a MHC class I immunodominant peptide from chicken ovalbumin, to the C-terminus of the outer membrane-associated Braun’s lipoprotein (Lpp) (Li et al., 2014). The manipulation of the EcN chromosome was carried out using a variation of the previously described CRISPR/Cas9 protocols (Qi et al., 2013; Zerbini et al., 2017), which allowed the in-frame insertion of the OVA sequence just upstream from the *lpp* stop codon (Figure S1). The expression of the Lpp-OVA fusion in EcN (EcN(*lpp-OVA*)) was confirmed by Western Blot analysis of total cell extract, using anti-OVA peptide antibodies (Figure S2).

Next, we assessed whether the oral administration of EcN(*lpp-OVA*) could elicit OVA-specific CD8^+^ T cells in the *lamina propria*. To this end, EcN(*lpp-OVA*) (10^9^ CFUs) was given by oral gavage to C57BL/6 mice three times at days 0, 3 and 6. One week after the last gavage mice were sacrificed and the presence of OVA-specific T cells in the *lamina propria* of small intestine was analyzed by flow cytometry (Figure 1B). As shown in Figure 1C the administration of EcN(*lpp-OVA*) elicited a significant fraction of OVA-specific CD8^+^ T cells in all treated mice (1.5-2.0% of total CD8^+^ T cells).

We then asked the question as to whether the administration of EcN(*lpp-OVA*) to C57BL/6 mice could influence the development of tumors when syngeneic OVA-B16F10 cells were injected subcutaneously. Mice (8 animals/group) were given either EcN or EcN(*lpp-OVA*) at days 0, 3 and 6 and one week after the last gavage all animals were challenged with 2.8×10^5^ OVA-B16F10 cells. Three additional gavages were administered, the first one the day after the challenge, and the other two at one-week intervals (Figure 1D). As shown in Figure 1E, the administration of EcN(*lpp-OVA*) delayed tumor development with statistical significance (P=0.0079).

### Oral administration of OMVs from E.coli strains engineered with different CD8^+^ epitopes induces epitope-specific T cell responses and inhibits tumor growth

Like all Gram-negative bacteria, EcN releases OMVs and since Lpp is a membrane-associated protein, Lpp-OVA fusion is expected to accumulate in the vesicular compartment. Therefore, the elicitation of the OVA-specific T cells observed after the oral administration of EcN(*lpp-OVA*) might be favored by the release of Lpp-OVA-decorated OMVs in the gut (Lpp-OVA-OMVs_EcN_). Thanks to their small size (30 - 300 nm in diameter), these vesicles should be efficiently taken up by mucosal APCs, thus promoting the local elicitation of OVA-specific T cells responses. Moreover, OMVs can cross the intestinal epithelium and reach the bloodstream, thus potentially eliciting T cells in other secondary lymphoid organs (Jones et al., 2020; Tulkens et al., 2020).

To test the possible contribution of microbiome-released OMVs in anti-tumor immunity, we first verified the production of OMVs by EcN(*lpp-OVA*) and the presence of the Lpp-OVA fusion protein in the vesicular compartment. EcN(*lpp-OVA*) was grown in liquid culture and at the end of the exponential growth the culture supernatant was subjected to ultracentrifugation and the pellet analyzed by SDS-PAGE. As shown in Figure S2, EcN(*lpp-OVA*) released OMVs, which carried a protein species of 10 kDa recognized by antibodies specific for the OVA peptide.

Lpp-OVA-OMVs_EcN_ were then purified from EcN(*lpp-OVA*) grown in a bioreactor and orally administered to C57BL/6 mice following the schedule reported in Figure 2A. One week after the last gavage, the animals were sacrificed and the presence of OVA-specific CD8^+^ T cells in the *lamina propria* was analyzed. As shown in Figure 2B, 4 to 6% of all recovered CD8^+^ T cells were OVA-specific.

**Figure 2.**
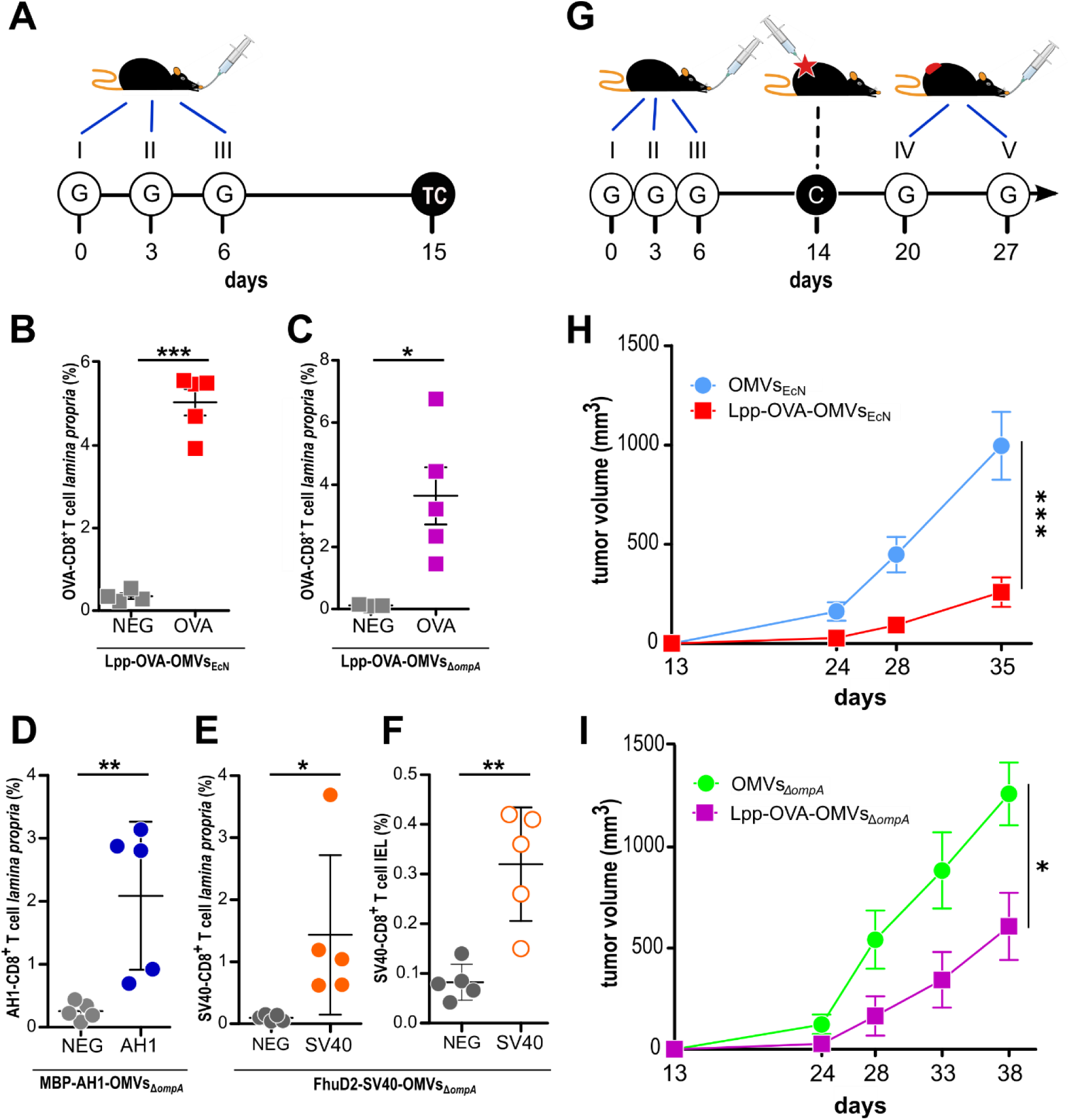
Effect of oral administration of OMVs on anti-tumor responses. (**A**) *Experimental protocol to analyze CD8*^*+*^*T cell responses after administration of OMVs engineered with T cell epitopes*. C57BL/6 mice were given three times 10 µg OMVs decorated with selected CD8^+^ T cell epitopes and one week after the last gavage epitope-specific T cells (TC) from *lamina propria* and/or intestinal epithelium (IEL) were analyzed by flow cytometry. (**B-F**) *Flow cytometry analysis of intestinal T cells*. Animals were given OMVs engineered with specific tumor epitopes as schematized in (A) and 1-2 × 10^6^ cells were isolated from the *lamina propria* (B-E) and intestinal epithelium (F) of the small intestine and the frequency of OVA-specific CD8^+^ T cells was measured by using epitope-specific Dextramer-PE. As negative control, the unrelated dextramer SSYSYSSL was used. (**G**) *Experimental protocol to study the tumor inhibitory activity of OMVs*. Mice were given OMVs by oral gavage (G) and were subsequently challenged with OVA-B16F10 tumor cells. Tumor growth was monitored for 23 days and during this period two additional gavages were administered. (**H-I**) *Analysis of OMV-mediated tumor inhibition*. C57BL/6 mice were treated with OMVs from wild type and OVA-expressing EcN and E. coli BL21 *ompA* as depicted in G and tumor volumes were measured over a period of 25 days. Statistical analysis was performed using Student’s t-test (two-tailed). ^*^ P ≤ 0.05; ^**^ P ≤ 0.01; ^***^ P ≤ 0.001.

Next, we asked the question as to whether Lpp-OVA-OMVs_EcN_ could protect C57BL/6 mice from the challenge with OVA-B16F10 cells. The vesicles were administered five times by oral gavage following the schedule reported in Figure 2G and tumor growth was followed for 22 days after challenging mice with tumor cells one week after the third oral gavage. As shown in Figure 2H, mice receiving Lpp-OVA-OMVs_EcN_ showed a substantial reduction in tumor growth when compared to mice that received “empty” OMVs (OMVs_EcN_) (P=0.0009).

We also investigated whether the tumor inhibitory activity of OMVs carrying the OVA CD8^+^ T cell epitope was restricted to vesicles released by EcN or rather vesicles from other *E. coli* strains could exert a similar function. To address this question, we fused the OVA epitope at the C-terminus of Lpp in the hyper-vesiculating *E. coli* BL21(DE3)Δ*ompA* strain (Fantappiè et al., 2014). The OMVs expressing the Lpp-OVA fusion (Figure S2) were purified from *E. coli* BL21(DE3)Δ*ompA*(*lpp-OVA*) culture and administered by oral gavage to C57BL/6 mice following the schedule previously described. As shown in Figure 2C, Lpp-OVA-OMVs_Δ*ompA*_ elicited OVA-specific CD8^+^ T cells in the *lamina propria* and protected mice from OVA-B16F10 challenge to an extent similar to what observed with Lpp-OVA-OMVs_EcN_ (P=0.0111) (Figure 2B).

Finally, we tested whether orally administered OMVs decorated with CD8^+^ T cell epitopes other than OVA could elicit epitope-specific T cells. To this aim, *E. coli* BL21(DE3)Δ*ompA* was engineered with two other epitopes, the AH1 peptide (SPSYVYHQF), derived from the gp70 envelope protein of the CT26 murine colon carcinoma cell line (Huang et al., 1996) and the SV40 epitope IV T antigen peptide VVYDFLKL, efficiently presented by MHC I molecules in C57BL/6 mice (Degl’Innocenti et al., 2005; Mylin et al., 2000). To drive the expression of the epitopes in the OMVs, two plasmids were generated, pET-MBP-AH1 and pET-FhuD2-SV40. pET-MBP-AH1 encodes the AH1 peptide fused to the C-terminus of Maltose Binding Protein (MBP), while pET-FhuD2-SV40 expresses the SV40 peptide fused to the C-terminus of FhuD2 (Irene et al., 2019). MBP-AH1-OMVs_Δ*ompA*_ and FhuD2-SV40-OMVs_Δ*ompA*_ were purified from *E. coli* BL21(DE3)*ΔompA*(pET-MBP-AH1) and *E. coli* BL21(DE3)Δ*ompA*(pET-FhuD2-SV40), respectively, and given three times, three days apart, by oral gavages to BALB/c and C57BL/6 mice, respectively (Figure 2A). One week after the last gavage, epitope-specific T cells were analyzed in the *lamina propria* and, in the case of SV40-C57BL/6, in the IELs population as well. As shown in Figure 2D-F, both OMVs elicited a relevant fraction of epitope-specific T cells, which could be detected both in the *lamina propria* and in the epithelium of the small intestine.

### The protective activity of EcN and OMVs engineered with a tumor epitope correlates with tumor infiltration of cancer-specific CD8^+^ T cells

The data described so far indicate that intestinal bacteria and OMVs have a broad capacity to induce CD8^+^ T cells against immunogenic epitopes in the gut and that the presence of these T cells correlate with tumor inhibition. Therefore, a plausible mechanism is that intestinal CD8^+^ T cells disseminate systemically and reach the tumors.

To support this mechanism, at the end of the challenge experiment depicted in Figure 1, two tumors from each group receiving either EcN or EcN(*lpp-OVA*) were surgically removed and the presence of total tumor infiltrating CD8^+^ T cells and of infiltrating OVA-specific CD8^+^ T cells was analyzed by flow cytometry. As shown in Figure 1F-G, a similar number of total CD8^+^ T cells was measured in tumors from animals that received EcN and EcN*(lpp-OVA*) (8.900 ± 500 CD8^+^ T cells/10^6^ live cells in both tumors). By contrast, the number of OVA-specific CD8^+^ T cells were two- and three-fold higher in tumors from mice treated with EcN(*lpp-OVA*) (9.2% of OVA-specific CD8^+^ T cells *vs* 3.88% of OVA-specific CD8^+^ T cells) (Figure 1G). The difference in the number of total and OVA-specific CD8^+^ T cells in tumors was confirmed by immunohistochemistry analysis of tumors from one animal per group stained with fluorescence-labelled OVA dextramers (CD8^+^ T cells_EcN*(lpp-OVA*)/_/CD8^+^ T cells_EcN_ = <u>1.05</u>; OVA-specific CD8^+^ T cells_EcN*(lpp-OVA*)/_/OVA-specific CD8^+^ T cells_EcN_ = <u>2.3</u>) (Figure 1G).

### Analysis of CD8^+^ T cell populations in the lamina propria and in the tumor by TCR sequencing

Having demonstrated that the oral administration of EcN(*lpp-OVA*) not only induced OVA-specific CD8^+^ T cells in the *lamina propria* but also increased the infiltration of OVA-specific CD8^+^ T cells in tumors, we tried to address the question as to whether the two OVA-specific CD8^+^ T cell populations might have a common origin. CD8^+^ T cells were isolated from both intestines and tumors of animals treated with either EcN or EcN(*lpp-OVA*) (Figure 3A) and their TCR sequences were compared. The analysis of the β chain sequences revealed a relatively high clonality of T cells in the tumors from mice that received the EcN gavages, while the T cell population in the *lamina propria* compartment of the same animals was much less diversified, as highlighted by the values of the Inverse Simpson Index as a measure of immune diversity (Figure 3B). The administration of EcN(*lpp-OVA*) appeared to reduce the diversification of T cells clones in the tumors. To evaluate the presence of recirculating and potentially tumor-specific T lymphocytes, GLIPH2 algorithm was employed to detect antigen-specific receptors on the basis of CDR3 similarity. The algorithm identified a specific CDR3 motif commonly shared in the *lamina propria* and in the tumors of EcN(*lpp-OVA*)-treated mice. The presence of this specific motif was statistically significant when compared to its occurrence in a mouse reference dataset (Figure S3). A deeper analysis of the TCR repertoire enabled the tracking of identical TCR sequences in different experimental groups. Interestingly, 11 TCR clonotypes were univocally present in the immune repertoire of T cells identified in the tumors and in the *lamina propria* of EcN(*lpp-OVA*)-treated mice. Five out of 11 TCR sequences could also be tracked in the tumors, but not in the *lamina propria*, of EcN-treated mice (Figure 3C).

**Figure 3.**
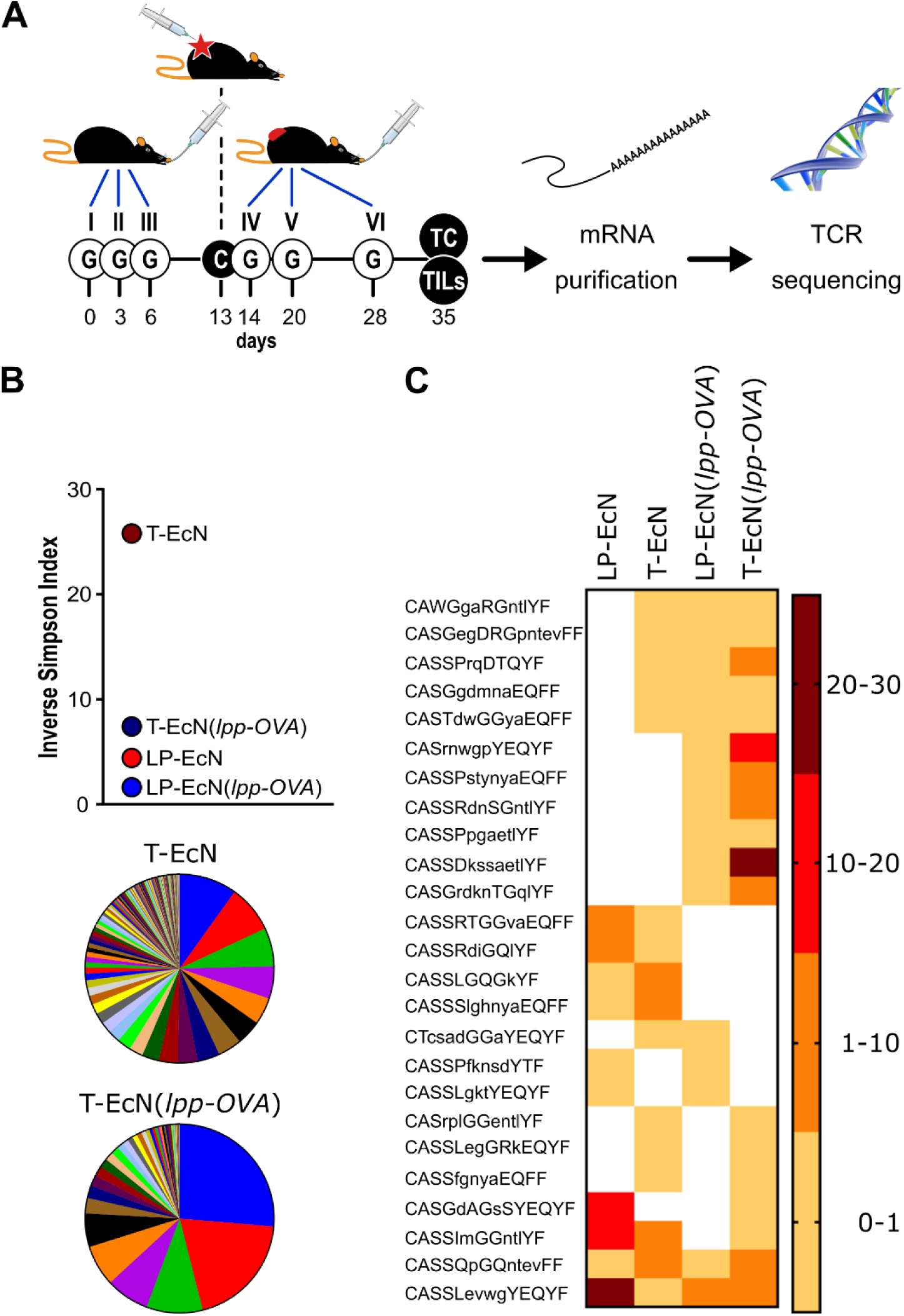
Analysis of TCR sequencing of CD8^+^ T cells from lamina propria and tumors. (**A**) Schematic representation of the experimental protocol. Mice (5 animals/group) were treated as depicted in Figure 3A and at the end of the experiment CD8^+^ T cells were collected from the *lamina propria* and tumor of each animal. After RNA extraction, the TCR β subunit was subjected to sequence analysis. (**B**) Analysis of TCR β subunit diversity using the Inverse Simpson Index. The analysis was carried out after pooling the TCR sequences of each group. The pie charts illustrate the different clonotypes identified in the tumors from EcN-treated mice (T-EcN) and from EcN(*lpp-OVA*)-treated mice (T-EcN(*lpp-OVA*). (**C**) Heat map showing the sharing of identical CDR3 amino acid sequences among different experimental groups. Color legend indicates the frequency of the clonotypes measured as relative sequence count. T: tumor; LP: *lamina propria*.

Overall, our data indicate that tumors from EcN-treated mice were infiltrated with a relevant number of T cell clones carrying TCRs identical to *lamina propria*-associated T cells (Figure 3C), which were likely elicited by the gut microbiome. However, these T cells were incapable to control tumor growth. The oral administration of the microbial species EcN(*lpp-OVA*) expressing a tumor-specific T cell epitope (OVA), resulted in the reorganization of the tumor infiltrating T cell population, with the concomitant enrichment of fewer T cell clones having TCRs identical to the TCRs of T cells found in the *lamina propria* and in the tumors of the same animals. Although not directly demonstrated, such T cells were probably those recognizing the OVA epitope and were originated at the intestinal site.

### Changes in microbiome composition after oral administration of EcN and OMV treatment

In addition, or as an alternative, to the direct role of OVA-specific CD8+ T cells, tumor inhibition could be mediated by modifications of the gut microbiome as a consequence of oral gavages. To exclude this possibility, we followed the microbiome composition via shotgun metagenomics in animals that were given oral gavages with EcN, EcN(*lpp-OVA*), OMVs_EcN_ and Lpp-OVA-OMVs_EcN_ and were subsequently challenged with OVA-B16F10 cells (Figure 4A, see Methods). The microbiome composition of mice after three subsequent oral administrations of all formulations displayed some changes as assessed by Bray-Curtis beta-diversity estimations (Figure 4B). Even though normal microbiome fluctuations are expected to occur longitudinally, the microbiome variations between T_0_ and T_1_ were less pronounced in mice receiving vesicles than bacteria. However, this effect was not due to the colonization of EcN since *E. coli* showed low relative abundances in all treatment groups (Figure S4). Importantly, the oral administration of either EcN or EcN(*lpp-OVA*) influenced the microbiome composition in a similar manner between T_0_ and T_1_ and between T_0_ and T_2_, calculated as Bray-Curtis dissimilarity (P = 0.84 and P = 0.69, respectively). The same effect was also observed after the gavage with OMVs_EcN_ or Lpp-OVA-OMVs_EcN_ (P = 1 and P = 0.57, respectively, Figure 4A). Therefore, the presence of Lpp-OVA in both EcN and OMVs did not induce more marked changes in microbiome. In fact, our data support the opposite: the administration of EcN and OMVs_EcN_ induced a significant change between T_0_ and T_1_(PERMANOVA P = 0.009 and P = 0.048, respectively) while no significant effect was seen in EcN(*lpp-OVA*) or Lpp-OVA-OMVs_EcN_ treated mice (PERMANOVA P = 0.105 and P = 0.057, respectively).

**Figure 4.**
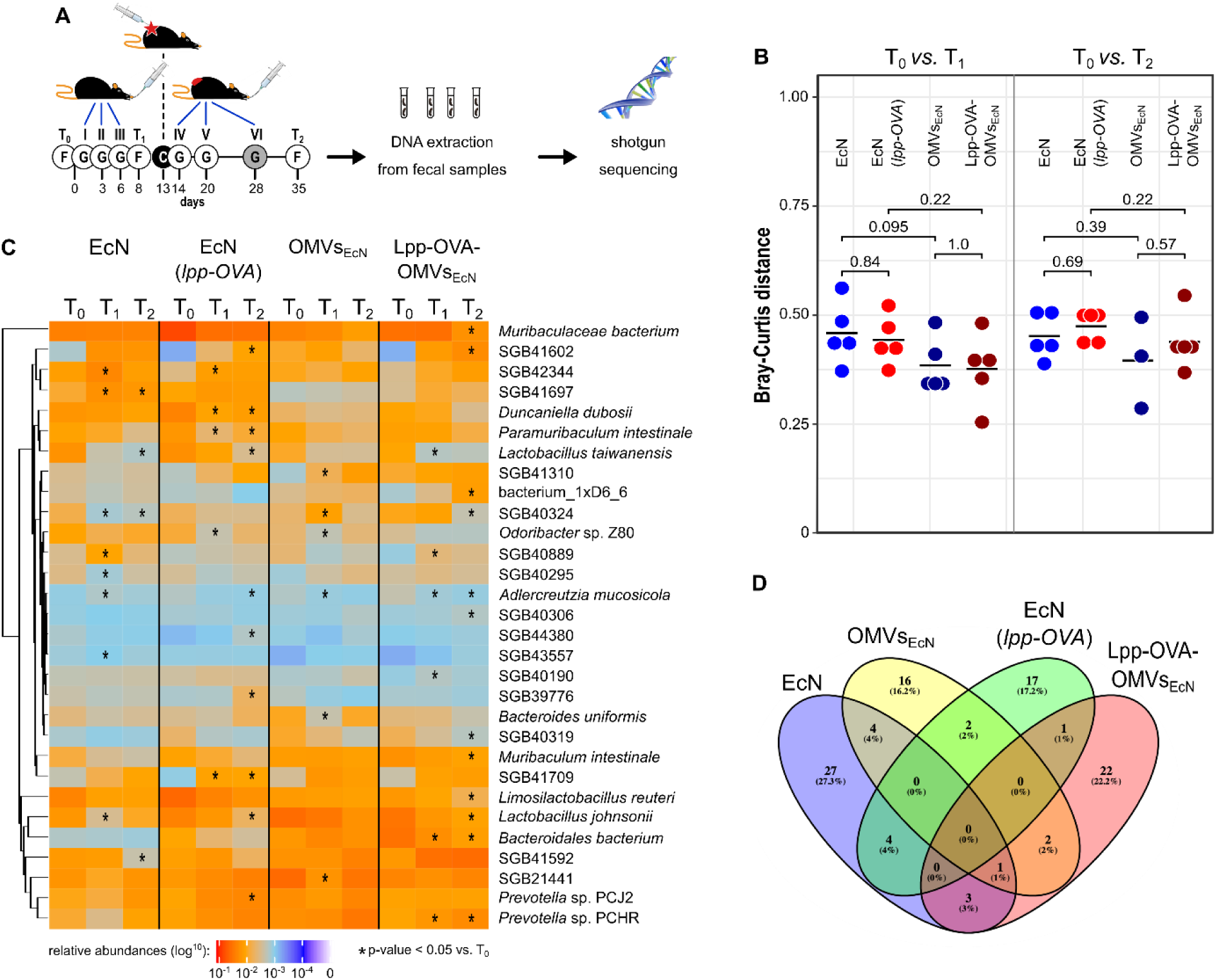
Effect of EcN, EcN(*lpp-OVA*), OMVs_EcN_ and Lpp-OVA-OMVs_EcN_ oral administration on gut microbiome composition. (**A**) *Schematic representation of the experimental procedure*. Mice were given three gavages (G) with either EcN, EcN(*lpp-OVA*), OMVs_EcN_ or Lpp-OVA-OMVs_EcN_. Mice were challenged with OVA-B16F10 cells (C) and then they received two (in the case of OMVs) or three additional oral administrations. Fecal samples were collected before the first gavage (T_0_), before the tumor challenge (T_1_) and at the end of the experiment (T_2_) and fecal DNA was subjected to shotgut sequencing. (**B**) Microbiome diversity in all mice within each treatment group at different time points (T_0_ *vs*. T_1_ and T_0_ *vs*. T_2_) estimated as Bray-Curtis distance. Statistical differences between group pairs were calculated through Wilcoxon Rank Sum Test. (**C**) Heatmap representing the relative abundance of the top 30 most abundant bacterial species that showed statistical variation within each group at different time points (T_0_ *vs*. T_1_ and T_0_ *vs*. T_2_) (* P ≤ 0.05). (**D**) Venn diagram showing the overlap of species with significant variations in relative abundance between T_0_ *vs*. T_1_ (Wilcoxon Rank Sum Test P ≤ 0.05).

While the microbiome change patterns are less clear at T_2_ (Figure 4B), potentially due to the tumor growth, this analysis provides evidence that the Lpp-OVA effect on tumor inhibition was not mediated by substantial overall changes of the microbiome. However, it could still be possible that specific taxa were directly or indirectly affected by the presence of the epitope. Considering the abundance of each microbial species we found several taxa that were significantly over- or under-represented (Figure 4C, Wilcoxon rank sum test, alpha 0.05). Nonetheless, little agreement was found on the panel of varying taxa among groups between T_0_ and T_1_ as shown by the heatmap in Figure 4C. In particular, among the groups that received a gavage with the OVA epitope, only one uncharacterized and yet-to-be cultivated species named SGB43006 was slightly significantly increasing at T_1_ compared to T_0_ (Figure 4D). SGB43006 abundance increased from 0.021% to 0.0395% in EcN(*lpp-OVA*) and from 0.0009% to 0.00946% in Lpp-OVA-OMVs_EcN_.

Taken together the data would indicate that the presence or absence of OVA in both EcN and OMVs was irrelevant with respect to the way the gut microbiome was perturbed by the animal treatment, thus strengthening the conclusion that the production of OVA-specific CD8^+^ T cells plays a direct role in tumor inhibition.

### OMVs engineered with a cancer epitope have a therapeutic effect against tumor growth

We finally asked the question as to whether OMVs decorated with cancer-specific epitopes could have a therapeutic effect on already implanted tumors. To address this question we challenged C57BL/6 mice with OVA-B16F10 cells and subsequently animals were given five oral gavages of either OMVs_Δ*ompA*_ or Lpp-OVA-OMVs_Δ*ompA*_ over a period of 21 days (Figure 5A). As shown in Figure 5B, tumor growth in mice treated with Lpp-OVA-OMVs_Δ*ompA*_ was delayed in a statistically significant manner with respect to animals receiving “empty” OMVs_Δ*ompA*_ (P=0.0031). Moreover, while four out of five mice treated with OMVs_Δ*ompA*_ reached a near to death status and therefore were sacrificed, none of the Lpp-OVA-OMVs_Δ*ompA*_-treated mice were sacrificed before the end of the experiment.

**Figure 5.**
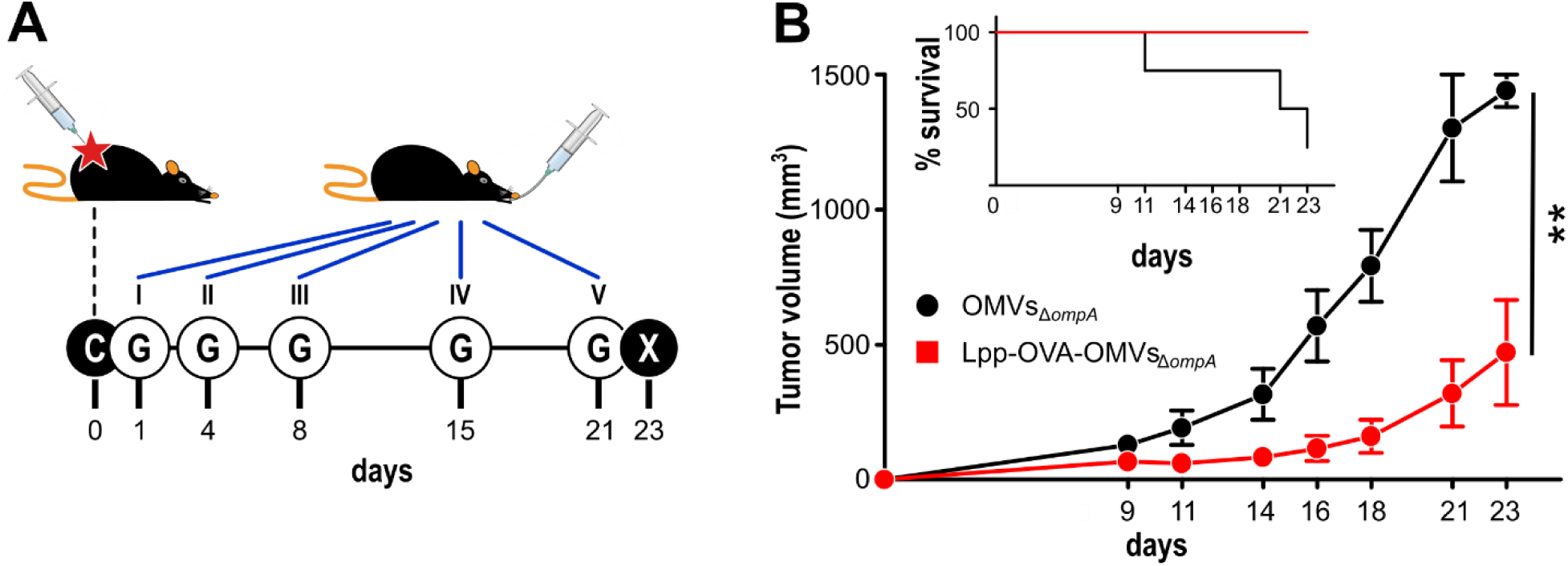
Therapeutic effect of Lpp-OVA-OMVs_Δ*ompA*_. (**A**) *Schematic representation of the therapeutic experimental protocol*. C57BL/6 mice (5 animals/group) were challenged with OVA-B16F10 tumor cells and subsequently treated with five oral administrations (G) of either OMVs_Δ*ompA*_ or Lpp-OVA-OMVs_Δ*ompA*_ (10 μg/dose) over a period of 23 days. (**B**) *Analysis of tumor growth inhibition*. Tumor volumes were measured at three day intervals and the average of tumor volumes from each group is plotted over time. The graph in the inlet shows the survival curve of each group (according to the authorized protocol, animals were sacrificed when tumors reached a volume of 1.500 mm^3^). Statistical analysis was performed using Student’s t-test (two-tailed). ^**^ P ≤ 0.01.

## Discussion

The underpinning mechanisms through which microbes influence cancer growth remain to be fully elucidated. The observations that patients responding and non-responding to checkpoint inhibitors can be stratified based on their microbiome composition (Routy et al., 2018b) and that the resistance to anti–PD-1 therapy in melanoma patients could be overcome by responder-derived fecal microbiota transplantation (FMT) (Davar et al., 2021) have prompted several laboratories to characterize the composition of the gut microbiome by 16S RNA/whole genome sequencing in search of microbial species with anti-tumor properties. *Enterococcus hirae, Bacteroides fragilis*, and *Akkermansia muciniphila* have been associated with favorable clinical outcome in cancer patients (Daillère et al., 2016; Rong et al., 2017; Routy et al., 2018b; Vétizou et al., 2015) and recently a pool of eleven bacteria has been proposed as a potential probiotic therapy (Tanoue et al., 2019). However, from all published data a “consensus” list of “anti-tumor bacterial species” remains to be unambiguously defined and it is safe to say that all studies converge to the conclusion that an important property of a “healthy, anti-tumor” microbiome is its diversity.

Regarding how microbiome exerts the anti-tumor effects, proposed mechanisms envisage the production of specific metabolites and/or the stimulation at the mucosal level of inflammatory cytokines which can work in a paracrine manner (Daillère et al., 2016; Pietrocola et al., 2016; Vétizou et al., 2015; Zitvogel et al., 2018). Also, gut-derived bacteria have been isolated from tumors, where they can promote pro- and anti-tumor effects (Geller et al., 2017; Hieken et al., 2016; Jin et al., 2019; Nejman et al., 2020; Pushalkar et al., 2018; Silva-Valenzuela et al., 2016). Finally, the “molecular mimicry” hypothesis has been proposed (Chai et al., 2017; Gil-Cruz et al., 2019; Perez-Muñoz et al., 2015; Rubio-Godoy et al., 2002; Vujanovic et al., 2007; Yang et al., 2014) holding that the anti-tumor effect of microbiome would be mediated by microbial-specific T cells, which accidentally recognize mutation-derived tumor neo-epitopes.

This work was conceptualized with the aim of demonstrating that cross-reactivity between the microbiome and cancer epitopes is involved in cancer immunity. We showed that when the human probiotic EcN expressing the OVA epitope was orally administered to C57BL/6 mice, high frequencies of OVA-specific T cells accumulated in the *lamina propria*. Moreover, oral delivery of EcN(*lpp-OVA*) reduced the growth of subcutaneous tumors in mice challenged with an OVA-B16F10 cell line. Importantly, tumor inhibition correlated with the elicitation of OVA-specific CD8^+^ T cells in the *lamina propria* and their infiltration in tumors, and our microbiome data supports the evidence that such T cells were the major players in the observed anti-tumor responses. As expected, the oral administration of both wild type EcN and EcN(*lpp-OVA*) modified the murine microbiome but the observed modifications were relatively modest and, importantly, not influenced by the expression of OVA in EcN. Even at the resolution of the shotgun sequencing, no specific single microbial species could be detected, which substantially differed from animals treated with wild type EcN and EcN(*lpp-OVA*). This would rule out the possibility that the anti-tumor activity of EcN(*lpp-OVA*) could be the result of an indirect effect acting on the alteration of other commensal species.

A question which remains to be fully addressed is the mechanism through the oral administration of EcN(*lpp-OVA*) promoted the enrichment of OVA-specific CD8^+^ T cells in the tumor environment. Our TCR sequencing data revealed the presence of T cell clones in the tumors and *lamina propria* T cells for EcN(*lpp-OVA*)-treated mice sharing identical TCRs, suggesting a common origin of these T cell populations. Therefore, the OVA-specific T cells induced by EcN(*lpp-OVA*) at the mucosal level could be disseminated systemically, eventually reaching tumor environment. An interesting observation emerging from our TCR sequencing data, is the high polyclonality of CD8^+^ T cells isolated from tumors of animals that received wild type EcN and a relevant number of TILs share TCRs identical to those found in the *lamina propria*. When animals are treated with EcN(*lpp-OVA*) TILs polyclonality is reduced. This would support the notion that tumors can be infiltrated with “exhausted” T cells which are incapable of effectively eliminated tumor cells. Thus, reducing the abundances of such T cells through the administration of microbiome species carrying properly selected cross-reactive epitopes could be a way to potentiate anti-tumor immunity.

OVA-specific CD8^+^ T cells were also found in tumors from EcN-treated animals, in line with the fact the all animals were challenged with an OVA-B16F10 cell line. Therefore, the possibility that the high frequency of OVA-specific CD8^+^ T cells observed in tumors of EcN(*lpp-OVA*)-treated mice was due to an amplification of locally induced T cells cannot be ruled out. As already pointed out, different microbial species have been isolated from animal and human tumors (Atreya and Turnbaugh, 2020) and such species often have intestinal origin. These intra-tumor bacteria could induce T cells (Kalaora et al., 2021), which eventually recognize tumor-specific cross-reacting epitopes.

An interesting piece of information emerging from our study is the contribution of OMVs to the elicitation of CD8^+^ T cells by intestinal bacteria. Oral delivery of OVA-OMVs derived from EcN(*lpp-OVA*) and *E. coli* BL21(DE3)Δ*ompA*(*lpp-OVA*) elicited high frequencies of OVA-specific CD8^+^ T cells and protected mice from the challenge with OVA-B16F10 cells. The capacity of OMVs to stimulate intestinal CD8^+^ T cells was not restricted to OVA as demonstrated by the isolation of CD8^+^ T cells after oral administration of MBP-AH1-OMVs_Δ*ompA*_ in BALB/c mice and FhuD2-SV40-OMVs_Δ*ompA*_ in C57BL/6 mice.

OMVs are fascinating organelles released by all Gram-negative bacteria with a plethora of biological functions such as intra- and inter-species cross talk and bacteria-host interactions (Kulp and Kuehn, 2010). Bacterial vesicles are known to be present in the intestine and they include both OMVs and vesicles produced by Gram-positive bacteria (Park et al., 2018). In the gastrointestinal tract, OMVs are believed to contribute to maintaining the intestinal microbial ecosystem and mediating the delivery of bacterial effector molecules to host cells to modulate their physiology. Shen et al. (Shen et al., 2012) showed that intestinal *Bacteroides fragilis* releases OMVs decorated with capsular polysaccharide (PSA). Dendritic cells sense OMV-associated PSA through TLR2 and stimulate the production of regulatory T cells, which protect from autoimmune and/or inflammatory diseases. Moreover, OMVs produced by the major human gut commensal bacterium *Bacteroides thetaiotaomicron* (Bt) have been shown to be acquired by intestinal epithelial cells via dynamin-dependent endocytosis followed by intracellular trafficking to endo-lysosomal vesicles. OMVs_Bt_ were also shown to transmigrate through epithelial cells via a paracellular route and to reach systemic tissues, leading to suggest that OMVs may act as a long-distance microbiota–host communication system (Jones et al., 2020). The capacity of bacterial vesicles to disseminate systemically in the host and their potential dual role in tumor development and inhibition has been recently reviewed (Chronopoulos and Kalluri, 2020).

Our work provides evidence of a broader OMV function, which extends to cancer immunity via MM. Thanks to their small size, OMVs can navigate in the gut and be easily internalized by intestinal DCs. The pan-proteome of OMVs from all gut bacteria has the potential to generate a large repertoire of T cells, which act as sentinels to eliminate host cells in which mutations eventually generate cross-reacting immunogenic epitopes. Such immunological role of OMVs, and potentially of all vesicles released by intestinal bacteria, would underline how inseparable evolution has made mammals and their microbiota and humans may be viewed as a single unit of evolutionary selection comprised of a host and its associated microbes (Rosenberg et al., 2009; Shen et al., 2012).

One last comment deserves the translational potential of our work. Our data show that the oral delivery of OMVs protects mice from tumor challenge both in the prophylactic and therapeutic modalities. This leads to the attractive hypothesis that OMVs engineered with cancer neo-epitopes could be exploited, in combination with other therapies such as checkpoint inhibitors, to potentiate the elicitation of cancer-specific T cell responses. The ease with which OMVs can be manipulated with multiple epitopes (Fantappiè et al., 2014, 2017; Grandi et al., 2017, 2018; Irene et al., 2019; Zanella et al., 2021) and can be purified from the culture supernatant, make the production of personalized oral cancer vaccines particularly attractive.

## Materials and Methods

### Bacterial strains, cell lines and mouse strains

DH5α, HK100 were used for cloning experiments. *Escherichia coli* Nissle 1917 (EcN) was isolated from the probiotic EcN^®^ (Cadigroup, Rome, Italy) and *E. coli* BL21(DE3)Δ*ompA* was produced in our laboratory (Fantappiè et al., 2014). *E. coli* strains were grown in LB at 37°C or 30°C in static or shaking conditions (200 rpm). When required, LB was supplemented with 50 µg/ml kanamycin, 25 µg/ml chloramphenicol, 0.2% L-arabinose and 5% sucrose. Stock preparations of *E. coli* strains in LB 20% glycerol were stored at -80°C.

OVA-B16F10 cell line, a B16F10 cell line transfected with a plasmid carrying a complete copy of chicken ovalbumin (OVA) cDNA and the Geneticin (G418) resistance gene, was kindly provided by Cristian Capasso and Prof. Vincenzo Cerullo from the Laboratory of Immunovirotherapy, Drug Research Program, Faculty of Pharmacy, University of Helsinki. OVA-B16F10 cell line was cultured in RPMI supplemented with 10% FBS, penicillin/streptomycin/L-glutamine and 5 mg/ml Geneticin™ (Gibco, Thermo Fisher Scientific, Waltham, MA, USA) and grown at 37°C in 5% CO_2_. C57BL/6 or BALB/c female 4-8 week old mice were purchased from Charles River Laboratories and kept and treated in accordance with the Italian policies on animal research at the animal facilities of Toscana Life Sciences, Siena, Italy and Department of Cellular, Computational and Integrative Biology (CIBIO) – University of Trento, Italy. Mice were caged in groups of 5/8 animals in ventilated cages. Mice within the same cage received the same treatment.

### Engineering EcN and E. coli BL21(DE3)ΔompA strains with CD8^+^ T cell epitopes

The pCRISPR-*lpp*-sgRNA plasmid, used for EcN mutagenesis, is a derivative of pCRISPR-*sacB* and codifies for a synthetic small guide RNA (sgRNA) (Zerbini et al., 2017). The *lpp*-sgRNA is composed by a 20 nt guide specific for *lpp*, a 42 nt Cas9-binding hairpin (Cas9 handle) and a 40 nt transcription terminator of *S. pyogenes* (Table S1) (Qi et al., 2013). For the construction of pCRISPR-*lpp*-sgRNA, a DNA fragment containing the *rrnB* T1 transcription terminator, the -10 and -35 consensus sequences of the J23119 promoter and the *lpp*-sgRNA chimera (Table S1) was synthesized by GeneArt (Thermo Fisher Scientific, Waltham, MA, USA) and cloned in pCRISPR-*sacB* using AvrII and XhoI restriction sites, thus replacing the gRNA cassette. The pCRISPR-*lpp*-gRNA plasmid, used for BL21(DE3)Δ*ompA* mutagenesis, is a derivative of pCRISPR-*sacB* in which a 30 nt DNA sequence coding for an *lpp*-gRNA guide specific for *lpp* is cloned using MB1360 and MB1361 oligonucleotides (Table S2) as described previously (Zerbini et al., 2017). Both pCRISPR-*lpp*-sgRNA and pCRISPR-*lpp*-gRNA contain a polylinker cloned into pCRISPR-*sacB* by PIPE-PCR with primers MB1346 and MB1347 and transformation in *E. coli* HK100 competent cells. The *lpp*-OVA donor DNA (dDNA) (Table S1) was chemically synthesized (GeneArt, Thermo Fisher Scientific, Waltham, MA, USA) with termini carrying the XhoI and NsiI restriction site sequences. It contains an EcN genomic region of 231 bp upstream from the *lpp* stop codon, followed by the OVA sequence flanked by restriction sites (NheI, Not and NdeI) and an EcN genomic region of 397 bp downstream from the *lpp* stop codon. In the arm upstream to OVA, the *lpp* TAC codon (Tyr), placed 9 bp upstream to the TAA stop codon, has been changed into TAT (Tyr) in order to eliminate the PAM sequence placed 5 bp upstream to the stop codon.

The construction of pET21-MBP-AH1 plasmid expressing the *E. coli* Maltose Binding protein (MBP) fused to three repeated copies of AH1 peptide linked by a Glycine–Serine (GS) spacers (Table S3) was obtained from pET21-MBP (Grandi et al., 2018) by ligating the AH1 DNA fragment carrying the BamHI/XhoI flags.

The pET21-FhuD2-SV40 plasmid carrying the *Staphylococcus aureus* Ferric hydroxamate receptor 2 (FhuD2) fused to one copy of SV40 IV peptide (Degl’Innocenti et al., 2005; Mylin et al., 2000), was assembled using the PIPE method (Klock and Lesley, 2009). Briefly, pET21-FhuD2 was linearized by PCR, using FhuD2-v-R and pET-V-F primers (Table S2). In parallel, the synthetic DNA encoding one copy of SV40 IV epitope (Table S3) was amplified by PCR with the forward FhuD2-SV40-F and the reverse FhuD2-SV40-R primers (Table S2). The PCR products were mixed together and used to transform *E. coli* HK100 strain. To confirm the correct gene fusions, plasmids were sequenced (Eurofins, Ebersberg, Germany, EU) and *E. coli* BL21(DE3)Δ*ompA* strain was transformed with pET21-MBP-AH1 and pET21-FhuD2-SV40 plasmids and the derived recombinant strain was used for the production of engineered MBP-AH1 and FhuD2-SV40 OMVs, respectively.

EcN(pCas9λred) and BL21(DE3)Δ*ompA*(pCas9λred) strains and competent cells were prepared as described previously (Zerbini et al., 2017). For genome engineering, 50 µl of EcN(pCas9λred) or BL21(DE3)Δ*ompA*(pCas9λred) competent cells were co-transformed with 100 ng of pCRISPR-*lpp*-sgRNA or pCRISPR-*lpp*-gRNA, respectively, and with 200 ng of dDNA. Cells were then incubated at 30°C (200 rpm) for 3 hours, plated on LB agar supplemented with chloramphenicol and kanamycin and incubated overnight at 37°C. The day after, single colonies were screened by colony PCR to identify clones carrying the OVA sequence insertion at the 3’-end of the *lpp* gene. Primers MB1336 and *lpp*2 (Table S2) were designed to anneal upstream and downstream the insertion site, thus generating amplicons of different lengths in the presence of *lpp-OVA* (609 nt) or wt *lpp* (549 nt). Positive clones were cured from pCRISPR-*sacB* derivative plasmids and from pCas9λred as described previously (Zerbini et al., 2017). The correctness of the *lpp-OVA* gene sequence was verified by sequencing using primers MB1336, *lpp*1, *lpp*2, MB1337, MB1390 (Table S2).

### OMV preparation

OMVs from EcN and EcN(*lpp-OVA*) were prepared growing the strains in an EZ control bioreactor (Applikon Biotechnology, Schiedam, Netherlands) as previously described (Zanella et al., 2021). Cultures were started at an OD_600_ of 0.1 and grown until the end of the exponential phase at 30°C, pH 6.8 (±0.2), dO2 > 30%, 280–500 rpm. OMVs were then purified and quantified as previously described (Zanella et al., 2021). BL21(DE3) *ompA* and BL21(DE3) *ompA*(*lpp-OVA*) were grown at 37°C and 180 rpm in LB medium (starting OD_600_ = 0.1) and when the cultures reached an OD_600_ value of 0.4-0.6 were maintained at 37°C under agitation for two additional hours. Finally, the purification of OMVs from BL21(DE3) *ompA*(pET-MBP-AH1) and BL21(DE3) *ompA*(pET-FhuD2-SV40) was carried out growing the cultures at OD_600_ = 0.5, adding 0.1 mM IPTG and continuing the incubation for 2 h at 37°C.

Culture supernatants were separated from biomass by centrifugation at 4000g for 20 minutes. After filtration through a 0.22-μm pore size filter (Millipore, Burlington, Massachusetts, USA), OMVs were isolated, concentrated and diafiltrated from the supernatants using Tangential Flow Filtration (TFF) with a Cytiva Äkta Flux system. OMVs were quantified using DC protein assay (Bio-Rad, Hercules, California. USA).

### OVA-Specific CD8^+^ T cells analysis in lamina propria and tumor tissue

For the analysis of T cells in the *lamina propria*, C57BL/6 or BALB/c mice were given bacteria (10^9^ CFUs) or OMVs (10 µg) by oral gavages at day 0, day 3 and 6 and at day 15 mice were sacrificed and small intestines were collected. In a first step, the intraepithelial lymphocytes (IELs) were dissociated from the mucosa by shaking the tissue in a pre-digestion solution, using the Lamina Propria Dissociation kit (Miltenyi Biotech, Bergisch Gladbach, Germany) according to the manufacturer’s instruction. Then the *lamina propria* tissue was treated enzymatically and mechanically dissociated into a single-cell suspension by using the gentleMACS™ Dissociators (Miltenyi Biotech, Bergisch Gladbach, Germany).

Tumor-infiltrating lymphocytes were isolated from subcutaneous OVA-B16F10 tumors as follows. Tumors (at least two tumors per group) were collected and minced into pieces of 1–2 mm of diameter using a sterile scalpel, filtered using a Cell Strainer 70 μm and transferred into 50-ml tubes. Then, the tumor tissue was enzymatically digested using the Tumor Dissociation kit (Miltenyi Biotech, Bergisch Gladbach, Germany) according to the manufacturer’s protocol and the gentleMACS™ Dissociators were used for the mechanical dissociation steps. After dissociation, the sample was passed through to a 30 μm filter to remove larger particles from the single-cell suspension.

At the end of the dissociation protocol, 1-2×10^6^ cells from *lamina propria* and tumors were incubated with 5 μl of OVA_257-264_ Dextramer-PE (SIINFEKL, IMMUDEX, Virum Denmark) for 10 minutes at room temperature in a 96-well plate. As negative control the unrelated dextramer SSYSYSSL was used (IMMUDEX, Virum Denmark). Then, cells were incubated with NearIRDead cell staining Kit (Thermo Fisher, Waltham, MA, USA) 20 minutes on ice in the dark. After two washes with PBS, samples were re-suspended in 25 μl of anti-mouse CD16/CD32-Fc/Block (BD Bioscience, San Jose, CA, USA), incubated 15 minutes on ice and then stained at RT in the dark for 20 minutes with the following mixture of fluorescent-labeled antibodies: CD3-APC (Biolegend, San Diego, CA), CD4-BV510 (Biolegend, San Diego, CA) and CD8a-PECF594 (BD Bioscience, San Jose, CA, USA). After two washes with PBS, cells were fixed with Cytofix (BD Bioscience, San Jose, CA, USA) for 20 minutes on ice, then washed twice and re-suspended in PBS. Samples were analyzed using a BD LSRII and the raw data were elaborated using *FlowJo* software. For the evaluation of the percentage of CD8^+^/OVA^+^ T cells in *lamina propria* and tumors the following gating strategy was applied. After selection of NearIRDead - cells (alive cells) and identification of FSC (forward scatter) and SSC (side scatter) morphology typical for the T cell population, only the SSW-cells (singlets) were selected and analyzed. This population was first separated in CD3^+^ and CD3^-^ cells and the CD3^+^ population was subsequently discriminated as CD4^+^ and CD8^+^ cells. Double positive CD8^+^/OVA^+^ T cells were finally selected.

### Mouse tumor models

Bacteria (10^9^ CFUs) and OMVs (10 μg) were given to C57BL/6 mice by oral gavage in 100 μl volume (PBS). C57BL/6 animals were subcutaneously challenged with 2.8×10^5^ OVA-B16F10 cells. Tumor growth was followed for at least 25 days after challenge and tumor volumes were determined with a caliper using the formula (A×B^2^)/2, where A is the largest and B the smallest diameter of the tumor. For the therapeutic protocol, mice were first challenged with 2.8×10^5^ OVA-B16F10 cells and subsequently treated with oral gavages (bacteria or OMVs) as described above. Tumor growth was monitored for at least 25 days. Statistical analysis (unpaired, two-tailed Student’s t-test) and graphs were processed using GraphPad Prism 5.03 software.

### Immunohistochemistry

Tumors from sacrificed mice were collected and maintained in RPMI on ice. Subsequently were covered with Tissue-Tek OCT compound and frozen with isopentane (VWR, Radnor, Pennsylvania, USA) kept in dry ice. 7-μm thick sections were cut from frozen OCT blocks, using Leica CM1950 Cryostat. Frozen sections were blocked with PBS 0.5% Bovine Serum Albumin (BSA, Sigma-Aldrich, St. Louis, MO, USA) followed by an overnight incubation at +4°C with anti-OVA_257-264_ Dextramer PE conjugate (SIINFEKL, IMMUDEX, Virum Denmark) diluted 1:30 in blocking solution. Then sections were fixed with PBS 2% paraformaldehyde (Sigma-Aldrich, St. Louis, MO, USA) and incubated with polyclonal Rabbit anti-PE (Abcam, Cambridge, UK) diluted 1:1.000 in PBS for 1 hour at room temperature. Subsequently, sections were incubated with goat anti-Rabbit Alexa-Fluor 488 conjugate (Molecular Probe, Waltham, MA, USA) diluted 1:500 in PBS, for 1 hour at room temperature. Sections were counterstained with 4′,6-Diamidino-2-phenylindole di-hydrochloride (DAPI, Sigma-Aldrich, St. Louis, MO, USA) diluted 1:3.000 in PBS, and then mounted with Dako Fluorescence mounting medium (Dako, Agilent Technologies, Santa Clara, CA, USA) and stored at +4°C until ready for image analysis. In order to confirm the specificity of the dextramer staining, a single immunofluorescence staining was performed to evaluate the effective presence of CD8^+^ cells. After an overnight air-drying step, frozen sections (see above) were fixed in Acetone (VWR, Radnor, Pennsylvania, USA) for 10 minutes, air-dried for 20 minutes and incubated with 5% goat serum in PBS (Sigma-Aldrich, St. Louis, MO, USA) to block non-specific reactions. Then, sections were incubated for 30 minutes at room temperature with a rat monoclonal antibody anti-Mouse CD8 (BD Pharmingen, BD Bioscience, San Jose, CA, USA) diluted 1:50 in blocking solution. Subsequently, sections were incubated for 1 hour at room temperature with goat anti-rat Alexa-Fluor 488 conjugate antibody (Molecular Probe, ThermoFisher Scientific Waltham, MA, USA) diluted 1:500 in PBS. Finally, sections were counterstained with DAPI, and mounted with Dako Fluorescence mounting medium as described above. Whole slide fluorescence images were acquired using Nanozoomer S60 automated slide scanner (Hamamatsu, Hamamatsu City, Japan). Positive cells were manually counted using NDP View2 Plus software (Hamamatsu, Hamamatsu City, Japan) on a total of 6 sections/tumor, covering a tissue depth of ∼300 μm. Total number of positive cells and area of tissue sections were measured, and cells density calculated as number of positive cells/mm^2^.

### TCR sequencing

At the end of the dissociation protocol (as described above), CD8^+^ cells from *lamina propria* and tumors were magnetically labeled with CD8 MicroBeads (Miltenyi Biotech, Bergisch Gladbach, Germany) according to the manufacturer’s manual. Then, the cell suspension was loaded onto a MACS^®^ Column, which was placed in the magnetic field of a MACS Separator. The magnetically labeled CD8^+^ T cells were retained within the column and eluted after removing the column from the magnetic field. At the end of separation, 5-8×10^3^ CD8^+^ T cells from *lamina propria* or 15-20×10^3^ CD8^+^ T cells from tumors were processed for the RNA extraction using the Arcturus^®^ PicoPure^®^ RNA Isolation Kit (Thermo Fisher, Waltham, MA, USA) according to the manufacturer’s protocol.

Complementarity determining region (CDR) 3 sequences of the TCR β chain were amplified by using a RACE approach (Bolotin et al., 2012). Samples were sequenced by using an Illumina MiSeq sequencer and CDR3 clonotypes identified using the MiXCR software (Bolotin et al., 2015). Sequences retrieved only once were excluded from the analysis. Normalized Shannon-Wiener index and Inverse Simpson index were calculated using the VDJtools package (PMID: 26606115). For comparing diversity indices between samples, original data were down-sampled to the size of the smallest dataset.

GLIPH2 algorithm (Huang et al., 2020) was employed to identify clusters of TCR sequences predicted to bind the same MHC-peptide. For this analysis, mMouse CD8 TCR set was selected as reference dataset and all amino acid were considered as interchangeable.

### Shotgun Whole Genome Sequencing of the gut microbiome

Bacteria or OMVs were administered to mice by oral gavage with 100 µl of PBS containing 10^9^ bacteria cells or 10µg of OMVs, respectively. Fecal samples were collected before the first gavage (T_0_), before tumor challenge (T_1_) and when animals were sacrificed (T_2_). Total DNA was purified from collected feces using the Quick-DNA Fecal/Soil Microbe Miniprep Kit (Zymo Research, Irvine, Canada, USA) according to the manufacturer’s instruction and subjected to shotgun sequencing. Sequencing libraries were prepared using the Illumina^®^ DNA Prep, (M) Tagmentation kit (Illumina, San Diego, California, USA), following the manufacturer’s guidelines. A cleaning step on the pool with 0.6× Agencourt AMPure XP beads was implemented. Sequencing was performed on a Novaseq600 S4 flowcell (Illumina, San Diego, California, USA) at the internal sequencing facility at University of Trento, Trento, Italy. Metagenomic shotgun sequences were quality filtered using trim galore discarding all reads of quality <20 and shorter than 75 nucleotides. Filtered reads were then aligned to the human genome (hg19) and the PhiX genome for human and contaminant DNA removal using Bowtie 2, v.2.2.8 (Langmead and Salzberg, 2012), yielding an average of 40 million bases in high-quality reads in each sample. Species-level microbial abundances were obtained through the bioBakery suite of tools using MetaPhlAn v.4.0 (Beghini et al., 2021) with default settings (database January 2021). Relative abundances at species level were analyzed in R. Beta diversity was calculated using the Bray-Curtis distance. Differential abundant species between T_0_/T_1_ and T_0_/T_2_ were discovered using a non-paired Wilcoxon test and selected based on a P < 0.05.

### SDS-PAGE and Western Blot analysis

In order to prepare total lysates, bacteria were grown in LB broth to an OD_600_ of 0.5, pelleted in a bench-top centrifuge and resuspended in Laemmli loading buffer to normalize cell density to a final OD_600_ of 10. OMVs were prepared as described above. 10 or 1 µl of total lysate and 10 or 1 µg of OMVs were separated on Criterion™ TGX Stain-Free™ any kD™ gel (Bio-Rad, Hercules, California, USA) for Coomassie staining or Western Blot, respectively, together with a protein marker (PM2610, SMOBIO Technology, Inc.). Gels were stained with ProBlue Safe Stain Coomassie (Giotto Biotech, Sesto Fiorentino Firenze, Italy, EU). For western blot, proteins were transferred onto nitrocellulose filters using iBlot™ gel transfer system (Invitrogen). Filters were blocked in PBS with 10% skimmed milk and 0.05% Tween for 45 min. and then incubated in a 1:1000 dilution of the required immune sera in PBS with 3% skimmed milk and 0.05% Tween for 60 minutes. Polyclonal antibodies against OVA or SV40 were obtained from GenScript (GenScript, Piscataway, New Jersey, USA) by immunizing rabbits with CGQLESIINFEKLTE or VVYDFLKC synthetic peptide, respectively. Filters were then washed 3 times in PBS-0.05% Tween, incubated in a 1:2000 dilution of peroxidase-conjugated anti-rabbit immunoglobulin (Dako, Santa Clara, California, USA) in PBS with 3% skimmed milk and 0.05% Tween for 45 min., washed 3 times in PBS-Tween and once in PBS. Before acquiring the signals, filters were treated with Amersham™ ECL Select™ Western Blot Detection reagent (GE Heathcare, Chicago, Illinois, USA).

## Acknowledgements

want to thank Cristian Capasso and Prof. Vincenzo Cerullo from the Laboratory of Immunovirotherapy (University of Helsinki) for kindly providing the OVA-B16F10 cell line. Furthermore, we gratefully thank Simona Tavarini and Chiara Sammicheli (Flow cytometry facility, GSK Siena, Italy) for flow cytometry acquisition and data analysis. We also highly appreciate the technical support of Carolina Fazio (CIO Preclinical Laboratories, TLS Siena, Italy) in *lamina propria* and tumor sample preparation and Daniela Fignani and Noemi Brusco (Fondazione Umberto Di Mario, Siena, Italy) for the technical support in IHC analysis.

## Author contributions

G.G. development of the overall concept, design of the research and writing of the manuscript. A.Gr. coordination of experimental activities. M.B., S.G., L.F. engineering bacteria with OVA epitope. A.Gr.; A.G., E.C., L.F. OMVs engineering with SV40 and AH1 epitopes. M.T., E.C., M.B., A.Gr., S.G., E.K. bacteria and OMVs preparations for oral administrations. E.C., A.Gr. S.T. T cell analysis in *lamina propria* and tumors. M.T., A.Gr., E.C., I.Z., E.K, L.C., S.V., E.B. animal experiments and tumor challenge models. G.L., E.C, G.S., A.Gr. immunohistochemistry experiments and imaging analysis. F.D. coordination of immunohistochemistry analysis. E.C., M.B., I.Z. DNA extraction for WGS analysis. N.S. management of microbiome and sequencing analysis, F.A. libraries preparation for WGS, M.D. WGS bioinformatics analysis. E.C., A.Gr. RNA extraction from CD8 T cell populations of *lamina propria* and tumors. E.R. TCR sequencing and data analysis. E.K., I.Z., M.B., S.G. figures editing. All: critical reading of the manuscript.

## References

Atreya, C.E., and Turnbaugh, P.J. (2020). Probing the tumor micro(b)environment. Science 368, 938–939.

Balachandran, V.P., Łuksza, M., Zhao, J.N., Makarov, V., Moral, J.A., Remark, R., Herbst, B., Askan, G., Bhanot, U., Senbabaoglu, Y., et al. (2017). Identification of unique neoantigen qualities in long-term survivors of pancreatic cancer. Nature 551, 512–516.

Beghini, F., McIver, L.J., Blanco-Míguez, A., Dubois, L., Asnicar, F., Maharjan, S., Mailyan, A., Manghi, P., Scholz, M., Thomas, A.M., et al. (2021). Integrating taxonomic, functional, and strain-level profiling of diverse microbial communities with bioBakery 3. Elife 10, e65088.

Bessell, C.A., Isser, A., Havel, J.J., Lee, S., Bell, D.R., Hickey, J.W., Chaisawangwong, W., Bieler, J.G., Srivastava, R., Kuo, F., et al. (2020). Commensal bacteria stimulate antitumor responses via T cell cross-reactivity. JCI Insight 5, e135597.

Bolotin, D.A., Mamedov, I.Z., Britanova, O. V., Zvyagin, I. V., Shagin, D., Ustyugova, S. V., Turchaninova, M.A., Lukyanov, S., Lebedev, Y.B., and Chudakov, D.M. (2012). Next generation sequencing for TCR repertoire profiling: Platform-specific features and correction algorithms. Eur. J. Immunol. 42, 3073–3083.

Bolotin, D.A., Poslavsky, S., Mitrophanov, I., Shugay, M., Mamedov, I.Z., Putintseva, E. V., and Chudakov, D.M. (2015). MiXCR: Software for comprehensive adaptive immunity profiling. Nat. Methods 12, 380–381.

Bresciani, A., Paul, S., Schommer, N., Dillon, M.B., Bancroft, T., Greenbaum, J., Sette, A., Nielsen, M., and Peters, B. (2016). T-cell recognition is shaped by epitope sequence conservation in the host proteome and microbiome. Immunology 148, 34–39.

Chai, J.N., Peng, Y., Rengarajan, S., Solomon, B.D., Ai, T.L., Shen, Z., Perry, J.S.A., Knoop, K.A., Tanoue, T., Narushima, S., et al. (2017). Helicobacter species are potent drivers of colonic T cell responses in homeostasis and inflammation. Sci. Immunol. 2, eaal5068.

Chronopoulos, A., and Kalluri, R. (2020). Emerging role of bacterial extracellular vesicles in cancer. Oncogene 39, 6951–6960.

Cook, D.P., Cunha, J.P.M.C.M., Martens, P.J., Sassi, G., Mancarella, F., Ventriglia, G., Sebastiani, G., Vanherwegen, A.S., Atkinson, M.A., Van Huynegem, K., et al. (2020). Intestinal delivery of proinsulin and IL-10 via Lactococcus lactis combined with low-dose anti-CD3 restores tolerance outside the window of acute type 1 diabetes diagnosis. Front. Immunol. 11, 1103.

Daillère, R., Vétizou, M., Waldschmitt, N., Yamazaki, T., Isnard, C., Poirier-Colame, V., Duong, C.P.M., Flament, C., Lepage, P., Roberti, M.P., et al. (2016). Enterococcus hirae and Barnesiella intestinihominis facilitate cyclophosphamide-induced therapeutic immunomodulatory effects. Immunity 45, 931–943.

Davar, D., Dzutsev, A.K., McCulloch, J.A., Rodrigues, R.R., Chauvin, J.M., Morrison, R.M., Deblasio, R.N., Menna, C., Ding, Q., Pagliano, O., et al. (2021). Fecal microbiota transplant overcomes resistance to anti-PD-1 therapy in melanoma patients. Science. 371, 595–602.

Degl’Innocenti, E., Grioni, M., Boni, A., Camporeale, A., Bertilaccio, M.T.S., Freschi, M., Monno, A., Arcelloni, C., Greenberg, N.M., and Bellone, M. (2005). Peripheral T cell tolerance occurs early during spontaneous prostate cancer development and can be rescued by dendritic cell immunization. Eur. J. Immunol. 35, 66–75.

Fantappiè, L., Santis, M. de, Chiarot, E., Carboni, F., Bensi, G., Jousson, O., Margarit, I., and Grandi, G. (2014). Antibody-mediated immunity induced by engineered Escherichia coli OMVs carrying heterologous antigens in their lumen. J. Extracell. Vesicles 3, 24015.

Fantappiè, L., Irene, C., De Santis, M., Armini, A., Gagliardi, A., Tomasi, M., Parri, M., Cafardi, V., Bonomi, S., Ganfini, L., et al. (2017). Some Gram-negative lipoproteins keep their surface topology when transplanted from one species to another and deliver foreign polypeptides to the bacterial surface. Mol. Cell. Proteomics 16, 1348–1364.

Fluckiger, A., Daillère, R., Sassi, M., Sixt, B.S., Liu, P., Loos, F., Richard, C., Rabu, C., Alou, M.T., Goubet, A.G., et al. (2020). Cross-reactivity between tumor MHC class I–restricted antigens and an enterococcal bacteriophage. Science. 369, 936–942.

Geller, L.T., Barzily-Rokni, M., Danino, T., Jonas, O.H., Shental, N., Nejman, D., Gavert, N., Zwang, Y., Cooper, Z.A., Shee, K., et al. (2017). Potential role of intratumor bacteria in mediating tumor resistance to the chemotherapeutic drug gemcitabine. Science. 357, 1156–1160.

Gil-Cruz, C., Perez-Shibayama, C., de Martin, A., Ronchi, F., van der Borght, K., Niederer, R., Onder, L., Lütge, M., Novkovic, M., Nindl, V., et al. (2019). Microbiota-derived peptide mimics drive lethal inflammatory cardiomyopathy. Science. 366, 881–886.

Grandi, A., Tomasi, M., Zanella, I., Ganfini, L., Caproni, E., Fantappiè, L., Irene, C., Frattini, L., Isaac, S.J., König, E., et al. (2017). Synergistic protective activity of tumor-specific epitopes engineered in bacterial Outer Membrane Vesicles. Front. Oncol. 7, 1–12.

Grandi, A., Fantappiè, L., Irene, C., Valensin, S., Tomasi, M., Stupia, S., Corbellari, R., Caproni, E., Zanella, I., Isaac, S.J., et al. (2018). Vaccination with a FAT1-derived B cell epitope combined with tumor-specific B and T cell epitopes elicits additive protection in cancer mouse models. Front. Oncol. 8, 481.

Hieken, T.J., Chen, J., Hoskin, T.L., Walther-Antonio, M., Johnson, S., Ramaker, S., Xiao, J., Radisky, D.C., Knutson, K.L., Kalari, K.R., et al. (2016). The microbiome of aseptically collected human breast tissue in benign and malignant disease. Sci. Rep. 6, 30751.

Huang, A.Y.C., Gulden, P.H., Woods, A.S., Thomas, M.C., Tong, C.D., Wang, W., Engelhard, V.H., Pasternack, G., Cotter, R., Hunt, D., et al. (1996). The immunodominant major histocompatibility complex class I-restricted antigen of a murine colon tumor derives from an endogenous retroviral gene product. Proc. Natl. Acad. Sci. U. S. A. 93, 9730–9735.

Huang, H., Wang, C., Rubelt, F., Scriba, T.J., and Davis, M.M. (2020). Analyzing the Mycobacterium tuberculosis immune response by T-cell receptor clustering with GLIPH2 and genome-wide antigen screening. Nat. Biotechnol. 38, 1194–1202.

Irene, C., Fantappiè, L., Caproni, E., Zerbini, F., Anesi, A., Tomasi, M., Zanella, I., Stupia, S., Prete, S., Valensin, S., et al. (2019). Bacterial outer membrane vesicles engineered with lipidated antigens as a platform for Staphylococcus aureus vaccine. Proc. Natl. Acad. Sci. U. S. A. 116, 21780–21788.

Jin, C., Lagoudas, G.K., Zhao, C., Blainey, P.C., Fox, J.G., Jacks, T., Jin, C., Lagoudas, G.K., Zhao, C., Bullman, S., et al. (2019). Commensal microbiota promote lung cancer development via gd T cells. Cell 176, 998–1013.

Jones, E.J., Booth, C., Fonseca, S., Parker, A., Cross, K., Miquel-Clopés, A., Hautefort, I., Mayer, U., Wileman, T., Stentz, R., et al. (2020). The uptake, trafficking, and biodistribution of Bacteroides thetaiotaomicron generated Outer Membrane Vesicles. Front. Microbiol. 11, 57.

Kalaora, S., Nagler, A., Nejman, D., Alon, M., Barbolin, C., Barnea, E., Ketelaars, S.L.C., Cheng, K., Vervier, K., Shental, N., et al. (2021). Identification of bacteria-derived HLA-bound peptides in melanoma. Nature 592, 138–143.

Klock, H.E., and Lesley, S.A. (2009). The Polymerase Incomplete Primer Extension (PIPE) method applied to high-throughput cloning and site-directed mutagenesis. Methods Mol. Biol. 498, 91–103.

Kulp, A., and Kuehn, M.J. (2010). Biological functions and biogenesis of secreted bacterial outer membrane vesicles. Annu. Rev. Microbiol. 64, 163–184.

Langmead, B., and Salzberg, S.L. (2012). Fast gapped-read alignment with Bowtie 2. Nat. Methods 9, 357–359.

Lasaro, M., Liu, Z., Bishar, R., Kelly, K., Chattopadhyay, S., Paul, S., Sokurenko, E., Zhu, J., and Goulian, M. (2014). Escherichia coli isolate for studying colonization of the mouse intestine and its application to two-component signaling knockouts. J. Bacteriol. 196, 1723–1732.

Li, G.W., Burkhardt, D., Gross, C., and Weissman, J.S. (2014). Quantifying absolute protein synthesis rates reveals principles underlying allocation of cellular resources. Cell 157, 624–635.

Mylin, L.M., Schell, T.D., Roberts, D., Epler, M., Boesteanu, A., Collins, E.J., Frelinger, J.A., Joyce, S., and Tevethia, S.S. (2000). Quantitation of CD8+ T-lymphocyte responses to multiple epitopes from Simian virus 40 (SV40) large T antigen in C57BL/6 mice immunized with SV40, SV40 T-antigen-transformed cells, or Vaccinia virus recombinants expressing full-length T antigen or epitope. J. Virol. 74, 6922–6934.

Nejman, D., Livyatan, I., Fuks, G., Gavert, N., Zwang, Y., Geller, L.T., Rotter-Maskowitz, A., Weiser, R., Mallel, G., Gigi, E., et al. (2020). The human tumor microbiome is composed of tumor type-specific intracellular bacteria. Science. 368, 973–980.

Park, K.S., Lee, J., Lee, C., Park, H.T., Kim, J.W., Kim, O.Y., Kim, S.R., Rådinger, M., Jung, H.Y., Park, J., et al. (2018). Sepsis-like systemic inflammation induced by nano-sized extracellular vesicles from feces. Front. Microbiol. 9, 1735.

Perez-Muñoz, M.E., Joglekar, P., Shen, Y.J., Chang, K.Y., and Peterson, D.A. (2015). Identification and phylogeny of the first T cell epitope identified from a human gut bacteroides species. PLoS One 10, e0144382.

Pietrocola, F., Pol, J., Vacchelli, E., Rao, S., Enot, D.P., Baracco, E.E., Levesque, S., Castoldi, F., Jacquelot, N., Yamazaki, T., et al. (2016). Caloric restriction mimetics enhance anticancer immunosurveillance. Cancer Cell 30, 147–160.

Pro, S.C., Lindestam Arlehamn, C.S., Dhanda, S.K., Carpenter, C., Lindvall, M., Faruqi, A.A., Santee, C.A., Renz, H., Sidney, J., Peters, B., et al. (2018). Microbiota epitope similarity either dampens or enhances the immunogenicity of disease-associated antigenic epitopes. PLoS One 13, e0196551.

Pushalkar, S., Hundeyin, M., Daley, D., Zambirinis, C.P., Kurz, E., Mishra, A., Mohan, N., Aykut, B., Usyk, M., Torres, L.E., et al. (2018). The pancreatic cancer microbiome promotes oncogenesis by induction of innate and adaptive immune suppression. Cancer Discov. 8, 403–416.

Qi, L.S., Larson, M.H., Gilbert, L.A., Doudna, J.A., Weissman, J.S., Arkin, A.P., and Lim, W.A. (2013). Repurposing CRISPR as an RNA-guided platform for sequence-specific control of gene expression. Cell 152, 1173–1183.

Rong, Y., Dong, Z., Hong, Z., Jin, Y., Zhang, W., Zhang, B., Mao, W., Kong, H., Wang, C., Yang, B., et al. (2017). Reactivity toward Bifidobacterium longum and Enterococcus hirae demonstrate robust CD8+ T cell response and better prognosis in HBV-related hepatocellular carcinoma. Exp. Cell Res. 358, 352–359.

Rosenberg, E., Sharon, G., and Zilber-Rosenberg, I. (2009). The hologenome theory of evolution contains Lamarckian aspects within a Darwinian framework. Environ. Microbiol. 11.

Routy, B., Gopalakrishnan, V., Daillère, R., Zitvogel, L., Wargo, J.A., and Kroemer, G. (2018a). The gut microbiota influences anticancer immunosurveillance and general health. Nat. Rev. Clin. Oncol. 15, 382–396.

Routy, B., Chatelier, E. Le, Derosa, L., Duong, C.P.M., Alou, M.T., Daillère, R., Fluckiger, A., Messaoudene, M., Rauber, C., Roberti, M.P., et al. (2018b). Gut microbiome influences efficacy of PD-1–based immunotherapy against epithelial tumors. Science. 359, 91–97.

Rubio-Godoy, V., Dutoit, V., Zhao, Y., Simon, R., Guillaume, P., Houghten, R., Romero, P., Cerottini, J.-C., Pinilla, C., and Valmori, D. (2002). Positional scanning-synthetic peptide library-based analysis of self- and pathogen-derived peptide cross-reactivity with tumor-reactive melan-A-specific CTL. J. Immunol. 169, 5696–5707.

Shen, Y., Torchia, M.L.G., Lawson, G.W., Karp, C.L., Ashwell, J.D., and Mazmanian, S.K. (2012). Outer membrane vesicles of a human commensal mediate immune regulation and disease protection. Cell Host Microbe 12, 509–520.

Silva-Valenzuela, C.A., Desai, P.T., Molina-Quiroz, R.C., Pezoa, D., Zhang, Y., Porwollik, S., Zhao, M., Hoffman, R.M., Contreras, I., Santiviago, C.A., et al. (2016). Solid tumors provide niche-specific conditions that lead to preferential growth of Salmonella. Oncotarget 7, 35169–35180.

Sivan, A., Corrales, L., Hubert, N., Williams, J.B., Aquino-Michaels, K., Earley, Z.M., Benyamin, F.W., Lei, Y.M., Jabri, B., Alegre, M.L., et al. (2015). Commensal Bifidobacterium promotes antitumor immunity and facilitates anti-PD-L1 efficacy. Science. 350, 1084–1089.

Sonnenborn, U. (2016). Escherichia coli strain Nissle 1917-from bench to bedside and back: History of a special Escherichia coli strain with probiotic properties. FEMS Microbiol. Lett. 363, fnw212.

Tanoue, T., Morita, S., Plichta, D.R., Skelly, A.N., Suda, W., Sugiura, Y., Narushima, S., Vlamakis, H., Motoo, I., Sugita, K., et al. (2019). A defined commensal consortium elicits CD8 T cells and anti-cancer immunity. Nature 565, 600–605.

Tulkens, J., Vergauwen, G., Van Deun, J., Geeurickx, E., Dhondt, B., Lippens, L., De Scheerder, M.A., Miinalainen, I., Rappu, P., De Geest, B.G., et al. (2020). Increased levels of systemic LPS-positive bacterial extracellular vesicles in patients with intestinal barrier dysfunction. Gut 69, 191–193.

Vétizou, M., Pitt, J.M., Daillère, R., Lepage, P., Waldschmitt, N., Flament, C., Rusakiewicz, S., Routy, B., Roberti, M.P., Duong, C.P.M., et al. (2015). Anticancer immunotherapy by CTLA-4 blockade relies on the gut microbiota. Science. 350, 1079–1084.

Vujanovic, L., Mandic, M., Olson, W.C., Kirkwood, J., and Storkus, W.J. (2007). A mycoplasma peptide elicits heteroclitic CD4+ T cell responses against tumor antigen MAGE-A6. Clin. Cancer Res. 13, 6796–6806.

Yang, Y., Torchinsky, M.B., Gobert, M., Xiong, H., Xu, M., Linehan, J.L., Alonzo, F., Ng, C., Chen, A., Lin, X., et al. (2014). Focused specificity of intestinal TH17 cells towards commensal bacterial antigens. Nature 510, 152–156.

Zanella, I., König, E., Tomasi, M., Gagliardi, A., Frattini, L., Fantappiè, L., Irene, C., Zerbini, F., Caproni, E., Isaac, S.J., et al. (2021). Proteome-minimized outer membrane vesicles from Escherichia coli as a generalized vaccine platform. J. Extracell. Vesicles 10, e12066.

Zerbini, F., Zanella, I., Fraccascia, D., König, E., Irene, C., Frattini, L.F., Tomasi, M., Fantappiè, L., Ganfini, L., Caproni, E., et al. (2017). Large scale validation of an efficient CRISPR/Cas-based multi gene editing protocol in Escherichia coli. Microb. Cell Fact. 16, 1–18.

Zitvogel, L., Ayyoub, M., Routy, B., and Kroemer, G. (2016). Microbiome and anticancer immunosurveillance. Cell 165, 276–287.

Zitvogel, L., Ma, Y., Raoult, D., Kroemer, G., and Gajewski, T.F. (2018). The microbiome in cancer immunotherapy: Diagnostic tools and therapeutic strategies. Science. 359, 1366–1370.

