## Supplementary material for "Outer Membrane Vesicles from the gut microbiome contribute to tumor immunity by eliciting cross-reactive T cells"

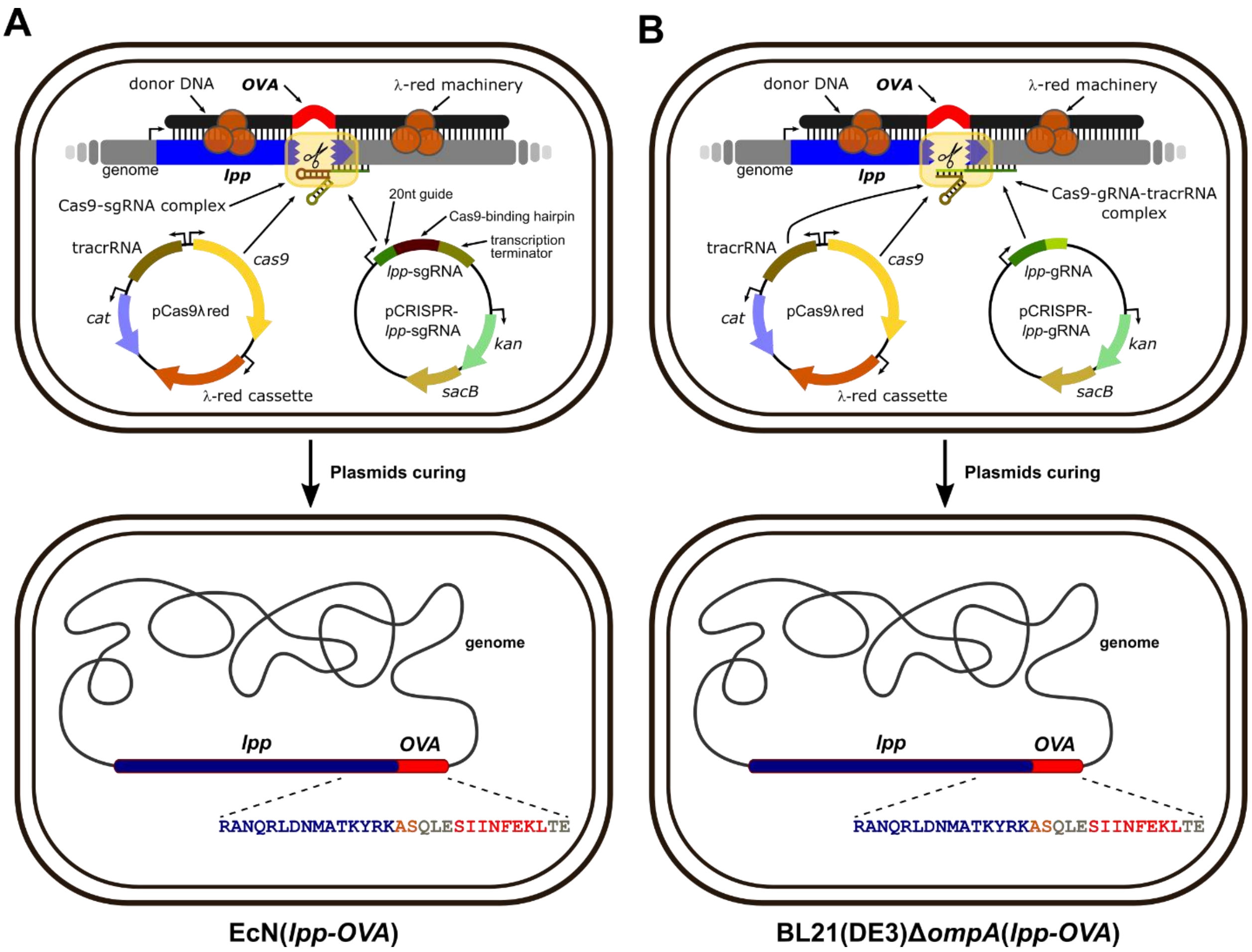


**Figure S1. Construction of *E. coli* strains engineered for Lpp-OVA fusion protein expression.** Schematic representations of the construction of EcN(*lpp-OVA*) (**A**) and *E. coli* BL21(DE3)Δ*ompA*(*lpp*-*OVA*) (**B**). EcN and BL21(DE3)Δ*ompA* were first transformed with the pCas9λred plasmid expressing the λ-Red proteins (orange circles), the Cas9 endonuclease (yellow rectangle) and the tracrRNA. Subsequently, EcN(pCas9λred) and BL21(DE3)Δ*ompA*(pCas9λred) were co-transformed with pCRISPR-*lpp*-sgRNA or pCRISPR-*lpp*-gRNA respectively and with a synthetic donor DNA (dDNA) carrying the OVA nucleotide sequence (in red) flanked by DNA fragments (in black) complementary to *lpp* (in blue) and the genome (in grey). In the system adopted for EcN(*lpp*-*OVA*) construction, the Cas9 enzyme is guided by a single RNA molecule (*lpp*-sgRNA) containing both the guide RNA and the Cas9-binding hairpin. Instead, in the system adopted for BL21(DE3)Δ*ompA*(*lpp*-*OVA*) construction, the Cas9 enzyme is guided by an RNA molecule deriving from the hybridization of the *lpp*-gRNA guide RNA with the tracrRNA coded from pCas9λred. Engineered clones with the *lpp*-*OVA* fusion gene were cured from plasmids. In the schematized resulting cells (bottom), the sequence of the C-terminal portion of the Lpp-OVA fusion protein is reported.

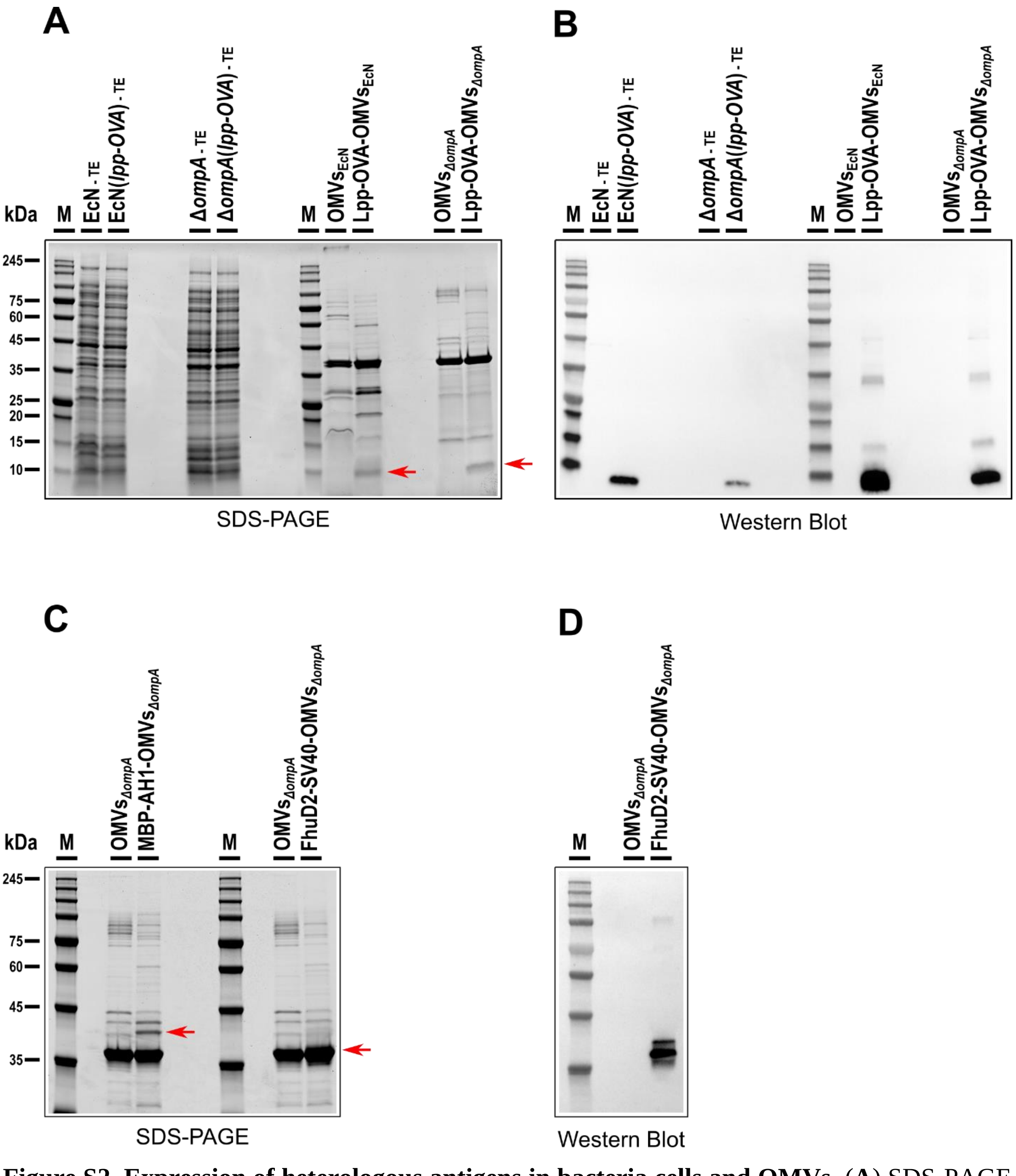


**Figure S2. Expression of heterologous antigens in bacteria cells and OMVs.** (**A**) SDS-PAGE and (**B**) Western Blot of total cell extracts (TE) and OMVs of *E. coli* strains EcN(*lpp-OVA*) and BL21(DE3)Δ*ompA*(*lpp-OVA*) (shortened to Δ*ompA*(*lpp-OVA*)) expressing Lpp-OVA fusion protein. For Western Blot anti-OVA polyclonal antibodies were used. (**C**) SDS-PAGE of OMVs derived from *E. coli* strains BL21(DE3)Δ*ompA*(pET-MBP-AH1) and BL21(DE3)Δ*ompA*(pET-

FhuD2-SV40) expressing MBP-AH1 and FhuD2-SV40 fusion proteins, respectively. (**D**) Western Blot of OMVs derived from *E. coli* strain BL21(DE3)Δ*ompA*(pET-FhuD2-SV40) expressing FhuD2-SV40 fusion protein using anti-SV40 polyclonal antibodies. EcN and BL21(DE3)Δ*ompA* (shortened to Δ*ompA*) were used as negative controls. Bands corresponding to recombinant antigens are indicated in SDS-PAGE panels by a red arrow. Protocols for sample preparation are described in supplementary material and methods.

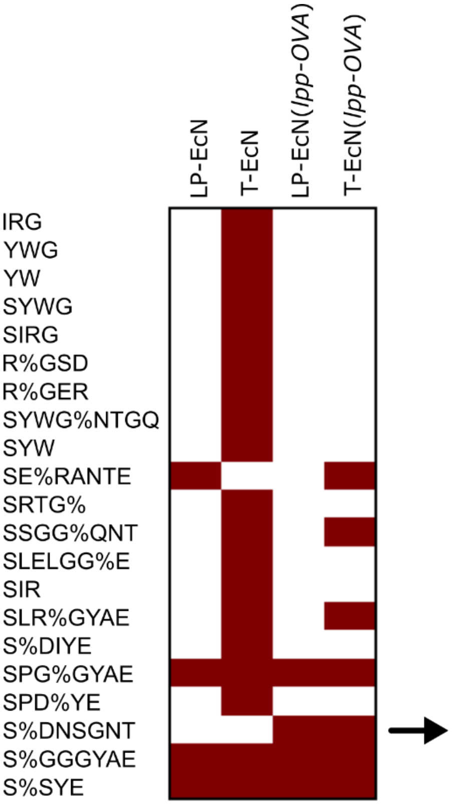


| Group | Sample | CDR3b | TRBV | TRBJ | Fisher score |
|---|---|---|---|---|---|
| global-S%DNSGNT | T-EcN(*lpp-OVA*) | CASSLdnSGntlYF | TRBV12-1 | TRBJ1-3 | 1.3e-03 |
| | LP-EcN(*lpp-OVA*) | CASSRdnSGntlYF | TRBV12-1 | TRBJ1-3 | |
| | T-EcN(*lpp-OVA*) | CASSRdnSGntlYF | TRBV12-1 | TRBJ1-3 | |

**Figure S3. Characterization of the TCR β chain repertoire**. (**A**) Normalized Shannon-Wiener index highlights the reduced diversity observed in the immune repertoire of $CD8^+$ T cells infiltrating the tumor of EcN(*lpp-OVA*)-treated mice compared to the EcN group. (**B**) Heat map showing the identification of CDR3 similarity motifs using the GLIPH2 algorithm. Fisher exact test is performed to estimate the bias of the pattern identified in the experimental datasets compared to a reference dataset. The Table shows the S%DNSDNG specificity group and the corresponding CDR3 amino acid sequences. The % indicates positions in the CDR3 sequences allowing variants. Amino acids in lower case are encoded by codon overlapping with N nucleotides added during the VDJ rearrangement process. TRBV: T cell receptor variable gene of the beta chain; TRBJ: T cell receptor joining gene of the beta chain. TCR sequencing results originated from the *lamina propria* of 4 EcN-treated mice, 3 EcN(*lpp-OVA*)-treated mice, and from the tumor of 5 EcN-treated mice and 2 EcN(*lpp-OVA*)-treated mice.

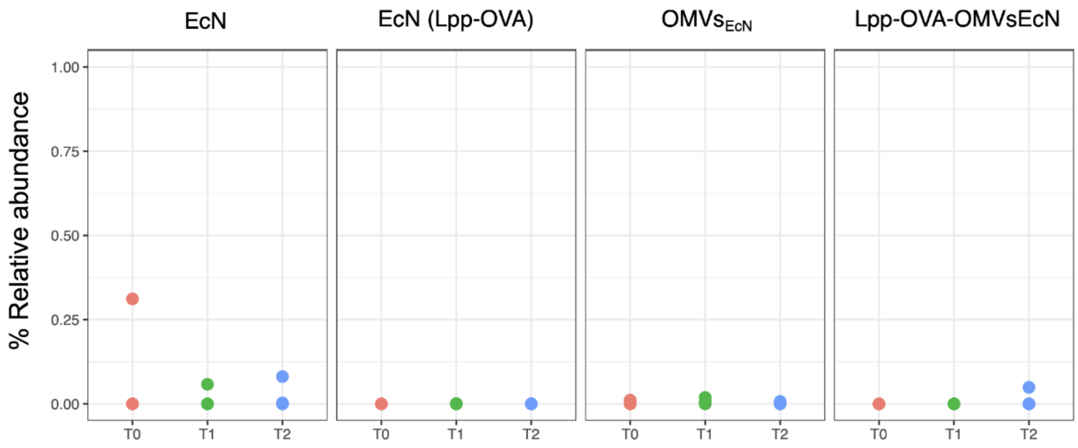


**Figure S4. Relative Abundance of Escherichia coli in fecal samples.** Plot representing the percentage of relative abundance of *E. coli* at $T_0$, $T_1$ and $T_2$ in each group treated as depicted in Figure 4A, calculated using MetaPhlAn v.4.0.

**Table S1. Sequences of guide and donor constructs used for mutagenesis.**

| Name | Nucleotide sequence |
|---|---|
| ***lpp*-sgRNA**[a] | CATATTCGTCTCCCTAGGTCTCAAATAAAACGAAAGGCTCAGTCGAAAGACTGGGC<br>CTTTCGTTTTATCTGTTGTTTGTCGGTGAACGCTCTCCTGAGTAGGACAAATACGCA<br>TCTGTGCGGTATTTCACACCGGATAAGCTGGATCCTTGACAGCTAGGTCAGTCCTA<br>GGTATAATACTAGT**CGGCATTTCACAGCATTAC**TGTTTTAGAGCTAGAAATAGCA<br>AGTTAAAATAAGGCTAGTCCGTTATCAACTTGAAAAAGTGGCACCGAGTCGGTGCT<br>TTTTTTGAATTCTCTAGAGTCGACCTGCAGAAGCTTAGATCTATTACCCTGTTATCC<br>CTACTCGAGTTC |
| ***lpp*-gRNA**[a] | **CATTTTTCACTTCACAGGTACTATTACTTG** |
| **dDNA**[b] | GGGACTCGAGAAAGCTACTAAACTGGTACTGGGCGCGGTAATCCTGGGTTCTACTC<br>TGCTGGCAGGTTGCTCCAGCAACGCTAAAATCGATCAGCTGTCTTCTGACGTTCAG<br>ACTCTGAACGCTAAAGTTGACCAGCTGAGCAACGACGTGAACGCAATGCGTTCCGA<br>CGTTCAGGCTGCTAAAGATGACGCAGCTCGTGCTAACCAGCGTCTGGACAACATGG<br>CTACTAAATATCGCAAGGCTAGC**CAGCTGGAAAGCATTATTAACTTTGAAAAAC**<br>**TGACCGAA**TAATGCGGCCGCCATATGTAGTACCTGTGAAGTGAAAAATGGCGCAC<br>ATTGTGCGCCATTTTTTTGCCTGCTATTTACCGCTACTGCGTCGCGCGTAACATATT<br>CCCTTGCTCTGGTTCCCCATTCTGCGCTGACTCTACTGAAGGCGCATTGCTGGCTGC<br>GGGAGTTGCTCCACTGCTCACCGCAACCGGATACCCTGCCCGACGATACAACGCTT<br>TATCGACTAACTTCTGATCTACAGCCTTATTGTCTTTAAATTGCGTAAAGCCTGCTG<br>GCAGCGTGTACGGCATTGTCTGAACGTTCTGCTGTTCTTCTGCCGATAGTGGTCGAT<br>GTACTTCAACATAACGCATCCCGTTAGGTTCCACGGAATATTTCACCGGTTCGTTGA<br>TCACTTTCACCGGTGTTCCCGTCCGCAATGCATTGG |

*lpp* guides[a] and OVA sequence[b] are in bold.

**Table S2. Oligonucleotides used in this study.**

| **Name** | **Nucleotide sequence** |
|---|---|
| MB1360 | AAACCATTTTTCACTTCACAGGTACTATTACTTGG |
| MB1361 | AAAACCAAGTAATAGTACCTGTGAAGTGAAAAATG |
| MB1346 | CATATGATGCATCCCGGGACGCGTGAAGACGAAAGGGCCTC |
| MB-1347 | ACGCGTCCCGGGATGCATCATATGACCTCGAGTCCCTATCAG |
| MB1336 | GATGAATCCGATGGAAGCATC |
| *lpp*2 | TCAGTAGAGTCAGCGCAG |
| *lpp*1 | GCTGTCTTCTGACGTTCAGAC |
| MB1337 | GATAAAGCGTTGTATCGTCGG |
| MB1390 | CTAGGACGTTACGCACTGC |
| FhuD2-v-R | TTTTGCAGCTTTAATTAATTTTTC |
| pET-v-F | CATCACCATCACCATCACGATTACA |
| FhuD2-SV40-F | TAATTAAAGCTGCAAAAGGCGGTGATAGCGTGGTG |
| FhuD2-SV40-R | GTGATGGTGATGTTATTACACCATCAGTTTCAGA |

**Table S3. Sequences of synthetic DNA encoding Gp70-AH1 and SV40 epitopes.**

| Name | Nucleotide sequence | Amino acid sequence |
|---|---|---|
| **Gp70-AH1 epitope** | ATTCAGGATCCAGCCCAAGCTATGTGTATCACCAATTCGGTTCCTCTCCATCCTATGTTTACCACCAGTTCGGCTCCTCGCCGAGCTATGTTTACCACCAGTTCTGACTCGAGTGAAT | SPSYVYHQFGSSPSYVYHQFGSSPSYVYHQF |
| **SV40 epitope** | GGCGGTGATAGCGTGGTGTATGATTTTCTGAAACTGATGGTG | GGDSVVYDFLKLMV |
